# Discovery of 5,5’-methylenedi-2,3-cresotic acid as a potent inhibitor of chemotactic activity of the HMGB1•CXCL12 heterocomplex using Virtual Screening and NMR validation

**DOI:** 10.1101/2020.08.25.266361

**Authors:** F. De Leo, G. Quilici, F. De Marchis, M. V. Mantonico, M. E. Bianchi, G. Musco

**Affiliations:** Biomolecular NMR Laboratory, IRCCS Ospedale San Raffaele, Division of Genetics and Cell Biology, Milan, Italy; Chromatin Dynamics Unit, Division of Genetics and Cell Biology, IRCCS Ospedale San Raffaele, Milan, Italy; Università Vita-Salute San Raffaele, Milan, Italy

**Keywords:** HMGB1, CXCL12, NMR, Virtual Screening, Chemotaxis, HMGB1/CXCL12 heterocomplex, HMGB1/CXCL12/CXCR4 axis

## Abstract

HMGB1 is a key molecule that both triggers and sustains inflammation following infection or injury, and is involved in a large number of pathologies, including cancer. HMGB1 participates to the recruitment of inflammatory cells forming a heterocomplex with the chemokine CXCL12 (HMGB1•CXCL12), herewith activating the G-protein coupled receptor CXCR4. Thus, identification of molecules that disrupt this heterocomplex can offer novel pharmacological opportunities to treat inflammation related diseases. To identify new HMGB1•CXCL12 inhibitors we have performed a study on the ligandability of the single HMG boxes of HMGB1 followed by a virtual screening campaign on both HMG boxes using Zbc Drugs and three different docking programs (Glide, AutoDock Vina, AutoDock 4.2.6). The best poses in terms of scoring functions, visual inspection and predicted ADME properties were further filtered according to a pharmacophore model based on known HMGB1 binders and clustered according to their structures. Eight compounds representative of the clusters were tested for HMGB1 binding by NMR. We identified 5,5’-methylenedi-2,3-cresotic acid (2a) as binder of both HMGB1 and CXCL12; 2a also targets the HMGB1•CXCL12 heterocomplex. In cell migration assays 2a inhibited the chemotactic activity of HMGB1•CXCL12 with IC50 in the subnanomolar range, the best documented up to now. These results pave the way for future structure activity relationship studies to optimize the pharmacological targeting of HMGB1•CXCL12 for anti-inflammatory purposes.

## Introduction

Cell recruitment is a fundamental event in the establishment of both acute and chronic inflammatory responses^1^. Chemokines and their receptors organize leukocyte trafficking and migration to the tissues both in healthy and pathological conditions^2^. Chemokines bind to and activate seven-transmembrane G-protein-coupled receptors (GPCRs), a receptor superfamily involved in many different diseases^3^. The different chemokines and their receptor are object of intense drug-discovery studies^4^. Increasing evidences show that chemokines usually require interactions with additional players that modulate the inflammation signaling cascade triggered by their receptors^5–7^; hence, the identification of molecules that disrupt the interactions of chemokines with their modulators might offer new and selective pharmacological opportunities against inflammation related diseases^8^. A prototype of functional synergic heterophilic interaction is represented by the heterocomplex formed by High Mobility Group Box 1 (HMGB1) and the chemokine CXCL12α, the 9 kDa ligand of the GPCR chemokine CXCR4 receptor^9^. The CXCL12/CXCR4 axis is crucial for chemotaxis, cell arrest, angiogenesis, cell survival and homing of hematopoietic progenitor cells in the bone marrow and their mobilization into the periphery both in physiological and pathological conditions^10–12^. The HMGB1•CXCL12 complex has been shown to trigger specific CXCR4 homodimer rearrangements^9^, ERK activation and calcium fluxes, along with enhanced CXCR4-dependent monocyte migration and tissue regeneration^13,14^. HMGB1 is a highly conserved, abundant non-histone nuclear protein (215 amino acids) that behaves as a DNA chaperone. Upon extracellular release, it acts as a DAMP (Damage Associated Molecular Pattern) and triggers cytokine-like proinflammatory and chemoattractant effects ^13,15^. HMGB1 is structurally organized in two independent L-shaped tandem domains (~80 amino acids each), BoxA and BoxB, followed by a 30 amino acid long acidic C-terminal tail^16,17^. Its modular organization, flexibility, different oxidation states^13,18^ along with its intracellular and extracellular localization confer to HMGB1 the ability to interact with many different partners, including nucleic acids, heparansulphates, lipopolysaccharides and proteins, thus exerting different functions spanning from architectural chromatin-binding activity^19^ to the modulation of innate immunity^20^. The cytokine-like function of HMGB1 upon inflammatory stimuli after infection or injury relies on the interaction of its reduced form with CXCL12, which promotes CXCR4-dependent recruitment of inflammatory cells to injured tissues^9^ and exacerbates the immune response in pathological conditions^5,21^. Thus, the disruption of the HMGB1•CXCL12 heterocomplex with small molecules might offer new and selective strategies against inflammation related diseases^22,23^. Interference with this heterophilic interaction is attractive but challenging for several reasons: i. the three-dimensional structure of the complex is still unknown, ii. the interaction surface is expected to be large and dynamic, thus difficult to be targeted by small molecules ^9,24,25^. Previous work has shown that it is possible to interfere with the pro-inflammatory properties of HMGB1 using small molecules including glycyrrhizin^26^, salicylic acid (SA) and its derivative amorfrutin^27^. Moreover, we have recently shown that diflunisal, an aspirin-like nonsteroidal anti-inflammatory drug, is able to selectively interfere *in vitro* and *in vivo* with the HMGB1/CXCL12/CXCR4 inflammatory axis by disrupting the interaction between HMGB1 and CXCL12^24^. This work has introduced the concept that the HMGB1•CXCL12 heterocomplex is a pharmacological target, herewith opening new perspectives for the rational design of novel inhibitors. While CXCL12 alone has been object of structure-based drug design studies and virtual screening campaigns aiming at inhibiting its direct activation effect on CXCR4^28,29^, systematic studies on the ligandability of HMGB1 and on the pharmacophoric requirements of HMGB1 ligands are still missing. This is in part ascribable to the L-shaped boxes of HMGB1, which do not appear to have clearly druggable pockets^30,31^, and to the absence of high resolution structures of HMGB1 in complex with known inhibitors.

In the present work we have explored the ligandability of HMGB1 and performed a structure-based virtual screening. The results were refined according to the pharmacophoric features of known HMGB1 ligands and tested for binding by NMR. This pipeline resulted in the identification of 5,5’-methylenedi-2,3-cresotic acid (**2a**) as a potent inhibitor of the chemotactic activity of HMGB1•CXCL12, with an IC_50_ in the subnanomolar range, and one of the best ligands identified up to now.

## Material and Methods

### Protein Structure Preparation

The structures of HMG boxes (BoxA, residues G3-Y77; BoxB, residues A93-G173) and CXCL12 (residues K1-K68) used for ligandability assessment, virtual screening (VS) and HADDOCK calculations were extracted from 2YRQ (first structure of the NMR bundle) and 4UAI, respectively^24^.

### *In silico* assessment of HMGB1 ligandability

Hot spots identification was performed using the FTMap computational map server (www.ftmap.bu.edu). The structures of BoxA and BoxB were uploaded into the FTMap server and ran according to instructions^32^. The server uses 16 organic molecules as probes and defines as consensus sites (CSs) the regions of the protein where several different probes clusters bind. The results were visually inspected using PyMol^33^.

Ligandability of HMGB1 boxes was further assessed using DoGSiteScorer (https://proteins.plus/). BoxA and BoxB structures were uploaded to the DoGSite server and scrutinized for binding sites, and their corresponding DrugScores were calculated according to the published protocol^34^. Pocket Size and DrugScores (the closest the score to 1, the higher the ligandability) were extracted for all identified sites and annotated to pocket numbers.

### Structure Based Virtual Screening

#### Library preparation

The Zbc Drugs subset (http://zinc12.docking.org/browse/subsets/special) containing 101746 biogenic drug-like compounds selected according to the Lipinski’s rule of five was chosen for virtual screening (VS) studies. The Zbc subset was prepared with the Ligand Preparation wizard available in the Schrödinger Suite, by adding hydrogens, minimizing and optimizing the molecules with OPLS_2005 force field. The ligands generated were ready for docking with GLIDE; for docking with AutoDock Vina^35^ and AutoDock 4.2.6^36^ they were converted to pdbqt-formatted file by in-house scripts. Ligands used to generate the training set for the pharmacophoric model (section 2.3.5) were prepared with the same procedure.

#### VS Docking studies

Protein-based VS studies were performed with three different programs: Glide (Schrödinger), AutoDock Vina^35^ and AutoDock 4.2.6^36^. Ligands were docked onto BoxA and BoxB.

##### GLIDE

BoxA and BoxB were prepared with the Protein Preparation Wizard available in the Schrödinger Suite: water molecules were removed, hydrogen atoms were added, the protonation states were adjusted according to neutral pH, finally the structures were minimized and optimized using OPLS2005 force field. For BoxA we generated a grid centered at 31.63, 27.3, 33.8 Å in correspondence of R23, the residue at the center of the experimentally validated binding pocket ^24^. The size of the inner box, that is the ligand diameter midpoint box, was left at the default values of 10 Å edges; the outer box, that is the box within which all the ligand atoms must be contained, was enlarged to 46 Å edges to allow ligands to find unusual or asymmetric binding modes in the active site. The grid for BoxB was centered on 27.22, 12.96, 7.84 Å in correspondence of R109 (the equivalent position of R23), the inner box was set at 10 Å edges, the outer box was set at 29 Å edges and the option to ‘dock ligands with length ≤19 Å’ was flagged. Glide High-Throughput Virtual Screening (HTVS) workflow was used for VS.

##### AutoDock VINA and AutoDock 4.2.6

BoxA and BoxB, prepared as described for Glide, were converted to pdbqt-formatted file. Charges and non-polar hydrogen atoms were added using the prepare_receptor4.py script from MGLTools. The binding site was defined by AutoGrid. The BoxA grid size consisted in 40 points in each direction; the grid point spacing was set at 0.458 Å and was centered on 31.6, 27.3, 30.70 on x, y, z axes, respectively. BoxB grid size was 40 Å in each direction, grid point spacing 0.531 Å and was centered on 20.5, 12.5, 3.0 on x, y, z axes, respectively. 2500000 max number of energy evaluations, 27000 max number of generations, 10 hybrid genetic algorithm-local search GA-LS runs were performed

#### Hits selection

The hit candidates emerging from Glide, AutoDock Vina and AutoDock 4.2.6 were selected according to energy ranking, distance filtering, visual inspection and absorption, distribution, metabolism, and excretion (ADME/Tox) descriptors through the QikProp module available in the Schrödinger Suite. In total, 34 properties and descriptors as defined in the Qikprop module were used to filter out the compounds outside the 95% of the value range reported for known drugs.

#### Pharmacophore model

The pharmacophore model (PhMOD) was generated using the automatic pharmacophore generation protocol in LigandScout3^37^ from Inte:Ligand. To generate the pharmacophore model we took advantage of a training set (TS) (Supplementary Figure S1A-B) composed by 7 known active HMGB1-interacting molecules from the literature, i.e. glycyrrhizin^26^, carbenoxolone^38^, salicylic acid (SA)^27^, amorfrutin^27^, acetyl-3-aminoethyl-SA Ac3AESA^27^, diflunisal^24^, inflachromene^39^ and 11 decoy HMGB1 ligands with an activity >50 μM as tested in-house in cell migration experiments (acetaminophen, ibuprofen, naproxen, nimesulide, ketoprofen, folic acid, tetrahydro methotrexate, cortisol, cortisone, prostaglandin E2) (Supplementary Figure S2). We superimposed the negatively charged moiety of the 3D-structures of the 7 active members of the TS and the following 4 pharmacophore features were generated: 5 hydrophobic moieties, 3 aromatic ring, 4 H-bonding acceptor and 1 H-bonding donor. The model was next refined maintaining only those features that previous structural studies on glycyrrhizin^26^ and diflunisal^24^ have indicated as relevant (Supplementary Figure S3A). The resulting PhMOD consisted of two H-bonding acceptor (HBA1 and HBA2) and two hydrophobics (H1 and H2). The tolerance sphere of HBA1-2 and H1 was increased by 0.15 Å and 0.3 Å, respectively, and H2 was set as optional feature to decrease the stringency of the PhMOD. The sensitivity of the PhMOD was measured by the ability to select among the TS the HMGB1 inhibitors in the first ranking positions, whereas specificity was measured by the ability of the PhMOD to identify HMGB1 inhibitors only. Sensitivity and specificity were represented in a receiver operating characteristic (ROC) curve to visualize the performance of the PhMOD (Supplementary Figure S3B). The ROC plot was calculated and visualized using LigandScout 3.02 and the AUC value (area under the ROC curve) was used to evaluate the ROC curve, with values between 0 (lowest) and 1 (highest) performance. The PhMOD was able to select active compounds significantly better than random (AUC=0.89) and with sufficient sensitivity and specificity for screening procedures (Supplementary Figure S3B). The Screening Perspective in LigandScout was used to filter the hit molecules resulting from the three VS programs against the three-dimensional PhMOD. Only molecules that fulfilled at least three out of four pharmacophore features were retained.

#### Hits clustering

The screened subsets were pooled together and clustered using Canvas 3.9 software (Schrödinger, LLC, New York, NY, 2020)^40^. The chemical fingerprint of each compound was first calculated using binary 2D linear fingerprint. Hierarchical clustering was then performed using the linear fingerprints with the average linkage method, which considers the average distance between all inter-cluster pairs. We obtained 20 clusters and analyzed only those with a compound population >10 molecules. We obtained 6 main clusters visualized using Hierarchical Clustering Dendrogram panel in Canvas. The clusters were numbered from the highest to the lowest populated: cluster 1 (148 molecules), cluster 2 (65 molecules), cluster 3 (47 molecules), cluster 4 (40 molecules), cluster 5 (25 molecules) and cluster 6 (20 molecules). The chemical scaffold representative of each cluster was calculated using CanvasMCS, the maximum common substructure facility. We set the fraction of molecules that must match the MCS as at least half the population of each cluster.

#### Protein expression and purifications

Recombinant HMGB1 constructs, including BoxA (residues 1–89), BoxB (residues 90–175), wildtype protein (Accession code P63158, residues 1-214), and mutant HMGB1 (R23A/R109A) in labelled and unlabeled forms were produced as described^24^. After expression and purification, proteins were dialyzed against NMR buffer, containing 20 mM phosphate buffer pH 7.3, 150 mM NaCl, 1 mM DTT.

Recombinant human CXCL12 (Accession code P48061, and residues M1-K69) was cloned into pET30a vector with restriction enzymes NdeI and BamHI. The resulting plasmid was transformed into *E.coli* BL21(DE3) and cells were grown at 37°C in Luria-Bertani medium. CXCL12 expression was induced by the addition of isopropyl-beta-D-thiolgalactopyranoside (IPTG) at a final concentration of 1 mM when cultures reached an optical density of 0.6-0.8 at 600 nm. Induced cultures were grown for additional 4 hours at 37°C, harvested by centrifugation at 6000 rpm and stored at −20°C until further processing. The cell pellets were resuspended in 20 ml buffer containing 50 mM TrisHCl (pH 8), 100 mM NaCl, 1 mM EDTA, 5 mM DTT, 0.1 mg DNAse, 0.1 mg RNAse and 5 mg lysozyme. The resuspended cells were lysed by sonication pulsed 1 second on and 1 second off for 2 minutes at 60% power. The inclusion bodies were washed twice with buffer containing 50 mM TrisHCl (pH 8), 100 mM NaCl, 1 mM EDTA, 5 mM DTT and finally with 50 mM TrisHCl (pH 8), 100 mM NaCl, 1 mM EDTA, 5 mM DTT, 0.1% Triton X100, followed by solubilization in buffer supplemented with 6M Guanidinium HCl and 50 mM HEPES pH 6.5 overnight at room temperature. The solubilized fraction was cleared by centrifugation at 18000 rpm at 4°C for 30 min and diluted dropwise into 250 ml refolding buffer containing 50 mM TrisHCl pH 7.4, 50 mM NaCl, 0.1 mM reduced glutathione, 0.1 mM oxidized glutathione and kept overnight at 4°C with stirring. Prior to chromatography, the protein solution was centrifuged at 18000 rpm for 30 min. CXCL12 was purified by cation-exchange chromatography using a SP Sepharose resin (SP Sepharose HP, GE Healthcare Bioscience AB, Uppsala, Sweden). The protein was washed with buffer A supplemented with 50 mM TrisHCl (pH 7.4) and 50 mM NaCl and eluted with buffer B containing 50 mM TrisHCl pH 7.4 and 1 M NaCl. CXCL12 was finally dialyzed against a buffer containing 20 mM phosphate buffer pH 6, 20 mM NaCl.

Protein concentrations were determined considering molar extinction coefficients at 280 nm of 21430 and 8700 M^−1^ cm^−1^ for HMGB1 and CXCL12, respectively.

Proteins used for cell-based assays were provided by HMGBiotech (Milan).

#### Reagents

ZINC4060879 (**1a**), ZINC2107574 (**1b**), ZINC12901682 (**1c**), ZINC4258914 (**3**), ZINC40310303 (**4a**), ZINC79188434 (**4b**), ZINC4215351 (**5**) were purchased from MCule (Budapest, Hungary). ZINC4028795/5,5’-methylenedi-2,3-cresotic acid (**2a**), ZINC57310 (**2b**) and ZINC988970/rosmarinic acid (**6**) were purchased from Sigma-Aldrich.

#### NMR Measurements

NMR spectra were recorded at 298 K on a Bruker Avance 600 MHz spectrometer (Karlsruhe, Germany) equipped with a triple-resonance TCI cryoprobe with an x, y, z-shielded pulsed-field gradient coil. Spectra were processed with Topspin^TM^ 3.2 (Bruker) and analyzed with CcpNmr Analysis 2.3^41^. ^1^H-^15^N-HSQC assignments of HMGB1 and CXCL12 were taken from the BMRB databank (HMGB1 accession numbers: 15148, 15149; CXCL12 accession number 16143)^42^ and confirmed *via* acquisition of 3D HNCA, CBCA(CO)NH experiments.

##### Saturation Transfer Difference and Water-Ligand Observed via Gradient Spectroscopy

STD and waterLOGSY experiments have been performed on 0.5 mM compounds (**1a**, **1b**, **2a**, **2b**, **4a**, **4b**, **5**, **6**) in the presence of 0.05 mM HMGB1 in NMR buffer. STD experiments were acquired using a pulse scheme (Bruker pulse sequence: stddiffesgp.3) with excitation sculpting with gradients for water suppression and spin-lock field to suppress protein signals. The spectra were acquired using 128 scans, a spectral width of 9600 Hz, 64K data points for acquisition. For protein saturation, a train of 60 Gaussian shaped pulses of 50 ms was applied, for a total saturation time of 3 s. Relaxation delay was set to 3 s. On-resonance irradiations were set at 10 ppm for **1a**, **1b**, **4b**, **5** and at 0 ppm for **2a**, **2b**, **4a**, **6**. Off-resonance was always set at 107 ppm. STD spectra were obtained by internal subtraction of the on-resonance spectrum from the off-resonance spectrum. WaterLOGSY experiments were acquired using a pulse scheme as described^43^ with excitation sculpting and flip-back for water suppression. The spectra were acquired using 128 scans, 32K data points for acquisition, mixing time was set to 1 s.

Ligand-based competition experiments were performed comparing STD and waterLOGSY spectra acquired on a sample containing 0.5 mM diflunisal and 0.05 mM CXCL12 with and without 1 mM **2a**.

##### Titrations

For NMR titrations, at each titration point a 2D water-flip-back ^1^H-^15^N-edited HSQC spectrum was acquired with 2048 (160) complex points for ^1^H (^15^N), respectively, apodised by 90° shifted squared (sine) window functions and zero filled to 256 points for indirect dimension. Titrations of **2a** and **6** into ^15^N HMGB1 were performed adding 0.5, 1, 2, 3, 5, 10 equivalents of ligand into the labelled protein. In order to minimize dilution and NMR signal loss, titrations were carried out by adding small aliquots of concentrated ligands (10 mM in 20 mM phosphate buffer, pH 7.3, 150 mM NaCl) to the ^15^N labelled protein samples (0.1 mM). Titration of **2a** on 0.1 mM ^15^N CXCL12 (20 mM phosphate buffer, pH 6, 20 mM NaCl) was performed adding 0.5, 1, 2, 3, 5, 10 equivalents of ligand to the labelled proteins using a 10 mM stock solution (20 mM phosphate buffer, pH 7.3, 150 mM NaCl). Assignment of the labelled proteins in the presence of the ligands (**2a**, **6**) was obtained following individual cross-peaks through the titration series. For each residue the weighted average of the ^1^H and ^15^N chemical shift perturbation (CSP) was calculated as CSP = [(Δδ_2_HN + Δδ_2_N/25)/2]_1/2_ ^44^. NMR-based antagonist-induced dissociation assays^45^ were performed by adding 0.5,1, 1.5, 2, 3, 5, 10 equivalents of **2a** on the ^15^N-HMGB1•CXCL12 heterocomplex (ratio 1:2).

##### Dissociation constant estimation

The apparent dissociation constants of **2a**-^15^N-HMGB1, **2a**-^15^BoxA, **2a**-^15^N-BoxB, **2a**-^15^N-CXCL12 and **6-**^15^N-HMGB1 interactions were estimated from least-squares fitting of CSPs as a function of total ligand concentration according to the equation:

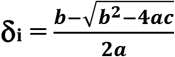

with a=(K_a_/δ_b_) [P_t_], b= 1+K_a_([L_ti_]+[P_t_]), and c=δ_b_K_a_[L_ti_], where δ_i_ is the absolute change in chemical shift for each titration point, [L_ti_] is the total ligand concentration at each titration point, [P_t_] is the total protein concentration, K_a_ =1/K_d_ is the association constant, and δ_b_ is the chemical shift of the resonance in the complex. The K_d_ of **2a**-^15^N-HMGB1, **2a**-^15^BoxA, **2a**-^15^N-BoxB, **2a**-^15^N-CXCL12 and **6-**^15^N-HMGB1 interactions were the average of 4, 3, 12, 3 and 4 residues, respectively, plus the standard deviation. For titrations of **2a** with HMGB1, based on previous data obtained on glycyrrhizin^26^ and diflunisal^24^, we assumed a stoichiometry of 2:1. Fitting the data of **2a** with the aforementioned equation yielded apparent K_d_ of 0.9, 1.2 and 1.1 mM for HMGB1, BoxA and BoxB respectively, thus supporting the presence of equivalent binding sites with similar apparent affinity (i.e., similar K_d_ values)^46^. K_d_ and δ_b_ were used as fitting parameters using the Xmgrace program (http://plasma-gate.weizmann.ac.il/Grace/).

^1^H resonance assignments of **2a** and **6** (1 mM dissolved in NMR buffer) were obtained analyzing 2D TOCSY (Total correlation spectroscopy) (mixing time 50 ms) and NOESY (Nuclear Overhauser effect spectroscopy) (mixing time 400ms) experiments.

Intermolecular nuclear Overhauser effect (nOes) between **2a** and BoxA were obtained from 3D ^13^C-NOESY-HSQC with no evolution on ^13^C dimension (2048 x 1 x 256 increments) experiments with ^15^N/^13^C filter in F1 (mixing time 200 ms); protein and ligand concentration were 0.8 mM and 1.6 mM, respectively, in D_2_O.

#### MST experiments

MST experiments were performed at 24°C on a NanoTemper^®^ Monolith NT.115 instrument with red filters, using 40% LED power and 60% MST power. Binding experiments were carried out using 6His-tagged HMGB1 and 6His-tagged CXCL12, non-covalently labelled with the NT647 fluorescence dye ^47^.

For binding assays **2a** was titrated (16-points) on 6His-tagged CXCL12 (MST buffer containing 20 mM phosphate buffer pH 7.3, 20 mM NaCl, 0.05% Tween). The ligand dilutions were generated as a 1:2 dilution of the stock solution using MST buffer; a constant amount of labelled proteins (50 nM) was added to all dilution steps. Maximum concentration of **2a** in the titration series was 5 mM. Complex samples were incubated for 15 minutes before loading into NanoTemper premium capillaries.

Competition experiments were carried out pre-forming a complex between labelled 6His-tagged HMGB1 (50 nM) and unlabeled CXCL12 (10 μM, *i.e*. 2 times the estimated K_d_)^48^. For 16-points titration series of **2a**, serial 1:2 dilutions of the **2a** stock solution were made into MST buffer, and a constant amount of pre-formed heterocomplex was added to all dilution steps. All samples were incubated for 15 minutes and centrifuged at 15,000 *g* for 10 minutes before measurements. Maximum concentration of **2a** in the titrations series was 5 mM. Addition of **2a** induced the recovery of the MST signal of HMGB1 towards the unbound state value.

For all MST experiments data points were the average of three measurements (error bars correspond to standard deviation). All data analyses were carried out using NanoTemper analysis software using the K_d_ model fitting for the binding assay and Hill model for competition experiment.

#### Data Driven Docking models

Molecular docking of **2a** on BoxA (residues G3-Y77), BoxB (A93-G173) (coordinates obtained as described in 2.1) were performed using the data-driven software HADDOCK 2.2^49,50^ following the classical three-stage procedure which includes: (1) randomization of orientations and rigid body minimization, (2) simulated annealing in torsion angle space, and (3) refinement in Cartesian space with explicit water. Ambiguous interaction restraints (AIRs) were defined as follows: residues with CSP > Avg + sd were used to define active residues, whose solvent accessible surface neighbors were set as passive (Supporting Table S1). In the case of CXCL12, only the residues located around the diflunisal binding site^24^ were set as active (Supporting Table S1), as STD competition experiments of **2a** in the presence of diflunisal demonstrated that they both compete for the same site. In the case of BoxA, intermolecular nOes were included as unambiguous restraints in the calculations only in the semi-flexible refinement stage, setting the maximum distance of the nOe H pairs to 5 Å (Supporting Table S1).

Optimized parameters for liquid simulation (OPLS) were used for the protein (protein-allhdg5-4 and protein-allhdg5-4-caro). The geometric coordinates and parameters for **2a** were calculated and optimized using the PRODRG server^51^. Calculations generated 1000, 1000, 500 structures for the rigid body docking (it0), the semi-flexible refinement (it1) and the explicit solvent refinement (water), respectively. The final 500 structures obtained after water refinement were scored with HADDOCK (HADDOCKscore = 1.0 E_vdW_ + 0.2 E_elec_ +1.0 E_desolv_ + 0.1 E_AIR_) for a weighted combination of van der Waals (vdW) and electrostatic energy terms (Lennard–Jones and Coulomb potentials), empirical desolvation term^52^ and ambiguous interaction restraint energy term, which reflects the accordance of the model to the input restraints.

HADDOCK models were clustered^53^ based on their interface root mean square deviation (rmsd), setting the cutoff and the minimum number of models in a cluster to 1.8 Å and 10 for the boxes, and 2.5 Å and 10 for CXCL12, respectively. Proteins were aligned and fitted on the backbone of active residues reported Supplementary Table S1. The rmsd of **2a** was calculated only on the heavy atoms of the entire scaffold.

To remove any bias of the cluster size on the cluster statistics, the final overall score of each cluster was calculated on the four lowest HADDOCK scores models in that cluster. For each protein the cluster with the best fitting relative to the experimentally-driven restraints (lowest number of violations) and the best HADDOCK score (cluster 1 for BoxA and BoxB, cluster 3 for CXCL12) was selected (Supplementary Figure S4A-C).

The analysis of the docking calculations was performed applying in-house python and tcl scripts. Molecular images were generated by PyMOL Molecular Graphics System, Version 2.0 Schrödinger, LLC and 3D Protein Imagining online server^54^.

#### Cell migration experiments

For fibroblast chemotaxis, modified Boyden chambers were used with filters (pore diameter 8 μm; Neuro Probe) coated with 50 μg/mL fibronectin (Roche). Mouse 3T3 cells (50,000 in 200 μL) were added to the upper chamber. Serum-free DMEM as negative control, HMGB1 and/or other molecules were added to the lower chamber at the indicated concentration, and then cells were left to migrate for 3 hours at 37°C. Cells were fixed with ethanol and stained with Giemsa Stain (Sigma), then non-migrating cells were removed with a cotton swab. All assays were done at least in biological triplicate. The migrated cells were acquired with Zeiss Imager M.2 microscope at 10x magnification, then evaluated with an automated counting program. All assays were done at least in biological triplicate and were repeated at least twice.

## Results

### *In silico* assessment of HMGB1 ligandability

We first searched for potential druggable hot spots on HMGB1 surfaces using the computational solvent mapping programs FTMap^32^ and Dogsite^34^. FTMap docks *in silico* small organic probes onto the surface of the protein and samples a large number of probe conformations, whereby contiguous areas of multiple probes clustering together might be considered as ligandable. FTMap identified 7 total consensus sites (CSs) on BoxA. CS2 CS3, CS5 and CS7 were within a distance of 8 Å, defining the highest density hot spot (39 probes), located at the interface of the two helices forming the short arm of the L-shaped fold (Figure 1A). Nine CSs were identified on BoxB; CS2 was the highest populated site, but did not have any other CS within 8 Å, thus could not be considered a hot spot. Conversely, CS1, CS8 and CS9, that were within 8 Å distance, created between the short helices of the BoxB the most populated hot spot (30 probes) (Figure 1B). The two hotspot regions identified on the two HMG boxes are characterized by small patches of partially solvent exposed hydrophobic residues including A16, F17, A19, F37 and F101, F102, F104, C105 in BoxA and BoxB, respectively. The ligandability of HMGB1 was further assessed by Dogsite, a web-based open-access algorithm that interrogates rigid protein structures for binding hotspots. The same regions identified by FTMap were the ones with the highest probability to be liganded, with a probability score DrugScore > 0.5 (DS = 0.58 for Box A and 0.73 for Box B) (Figure 1C-D). Collectively, both FTMap and Dogsite suggest that both HMG boxes are ligandable.

**Figure 1.**
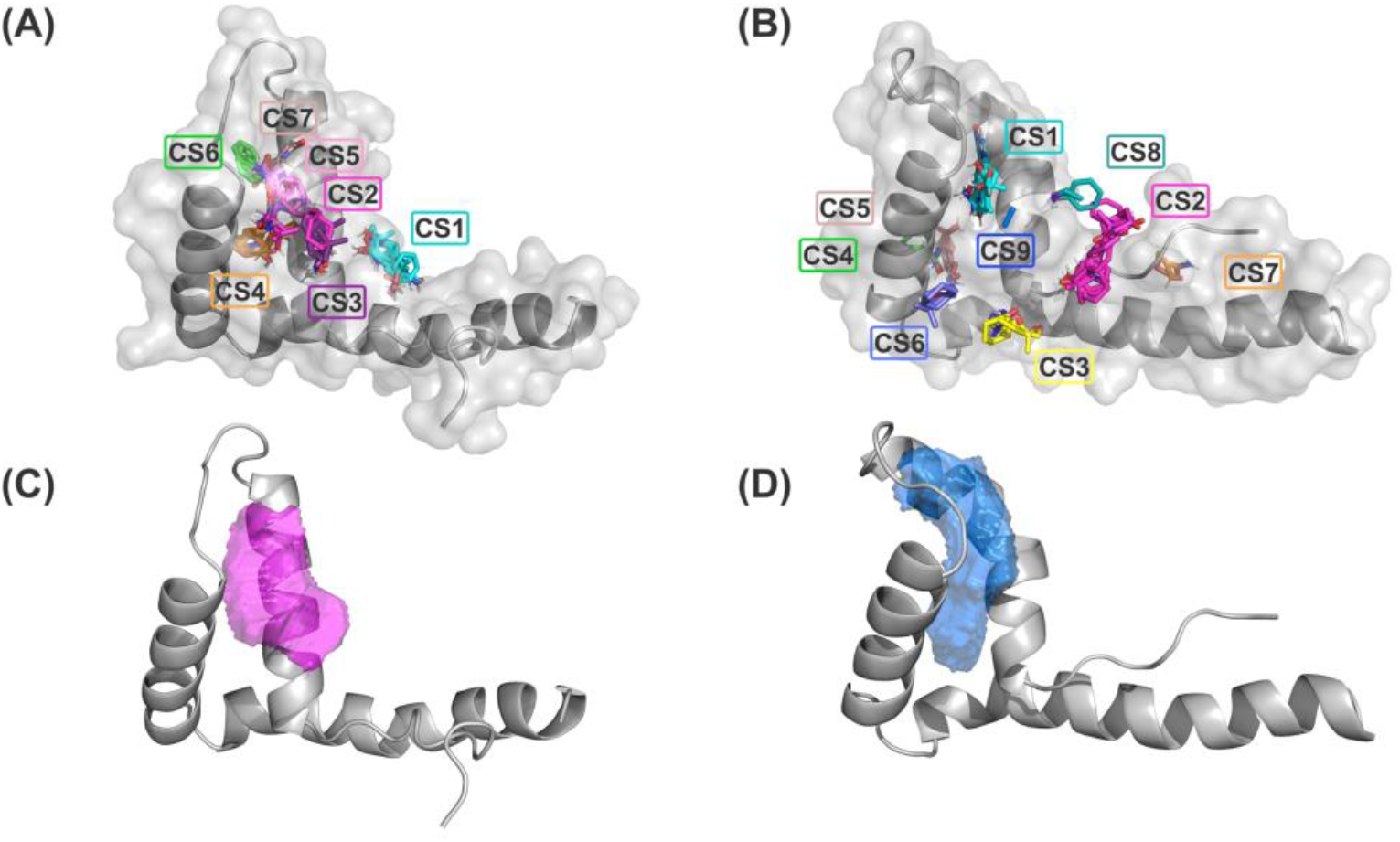
Ligandability assessment of HMG boxes by FTMap and Dogsite. FTMap predicts 7 Consensus Sites (CS) on BoxA (A) and 9 CS on BoxB (B). The probes (in sticks) populating the different CSs are colored accordingly. HMG boxes are represented in grey (cartoon and surface). The ligand binding pocket volumes with the highest DrugScores as predicted by Dogsite are represented by magenta and blue mesh surfaces on BoxA (C) and BoxB (D), respectively. HMG boxes are represented in grey (cartoon).

### Virtual Screening

We next screened the Zbc Drugs subset (101746 compounds) on BoxA and BoxB using three different docking programs, i.e. Glide, AutoDock Vina and AutoDock 4.2.6, and their results were combined (Figure 2). The three programs rely on distinct empirical scoring functions^55,56^ and differently weighted electrostatic and hydrophobic interactions. Thus, merging of their outputs was expected to mitigate the possible biases towards hydrophobic or electrostatic contributions characteristic for AutoDock Vina, AutoDock 4.2.6 and Glide, respectively.

**Figure 2.**
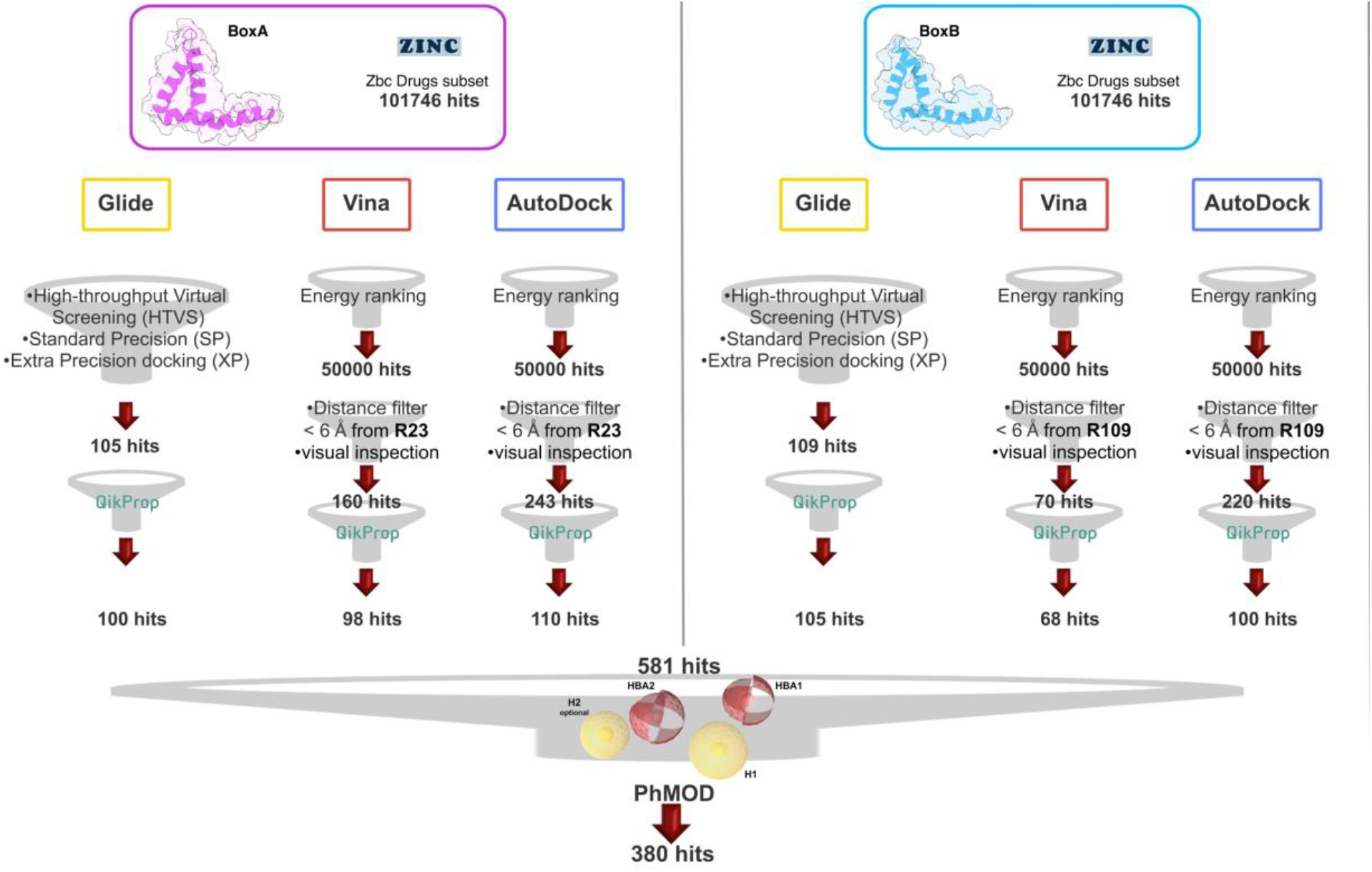
VS workflow. Scheme of the filtering steps adopted during the docking procedures computed with AutoDock Vina, and AutoDock 4.2.6 on BoxA (left) and BoxB (right). The filtering criteria and the number of hits selected at each filtering step are explicitly indicated. The red and the yellow spheres of the PhMOD represent the H-bonding acceptors (HBA1 and HBA2) and the hydrophobic (H1 and H2) moieties, respectively

The Virtual Screening (VS) workflow implemented in Glide (High Throughput Virtual Screening, HTVS; single precision, SP; extra precision XP docking) retrieved 105 compounds for BoxA and 109 for BoxB, ranked according to the scoring function assigned to the poses. For this subset the ADME descriptors were predicted, and molecules with ADME values outside the 95% range calculated for known drugs were discarded, yielding 100 and 105 hits for BoxA and for BoxB, respectively. Filtering according to ADME descriptors excluded molecules with low octanol/water partition coefficient (logP_oct/wat_), low predicted binding affinity to human serum albumin, high dipole moment, high electron affinity and low brain/blood partition coefficient (logBB), all properties in line with the known bias of Glide towards charged and hydrophilic molecules^57^.

For the VS performed with AutoDock Vina and AutoDock 4.2.6 the first 50000 top ranked poses were filtered retaining the ones in which the ligand was at a distance ≤ 6 Å from R23 in BoxA and R109 in BoxB. These arginines are crucial for ligand binding both for glygcyrrhizin^26^ and diflunisal^24^. Next, visual inspection of the AutoDock Vina/AutoDock 4.2.6 poses yielded 160/243 molecules for BoxA and 70/220 molecules for BoxB (Figure 2).

The molecules were next filtered according to ADME descriptors, retaining 98/110 molecules for BoxA and 68/100 molecules for BoxB (Figure 2). The characteristics of the discarded molecules reflected the bias of both AutoDock Vina/AutoDock 4.2.6 towards the selection of hydrophobic hits. In fact they were predicted to be or to generate metabolites with a high aromatic component of the solvent accessible surface area (SASA), low aqueous solubility (log S), and low values of H-bond acceptor moieties, hydrophilic SASA, dipole moment, water/gas partition coefficient.

Overall, VS using the three different programs yielded three sets for BoxA and three for BoxB for a total of 581 possible hits. This merged set was further interrogated using a pharmacophore model (PhMOD) generated considering known active HMGB1 inhibitors as described in Materials and Methods. The PhMOD consisted in two H-bonding acceptors (HBA1, HBA2) and two hydrophobic moieties (H1, H2) (Figure 2). Pharmacophore screening of the 581 possible hits resulted in 380 structures fulfilling at least 3 out of the 4 pharmacophoric requirements. These structures were then subjected to fingerprint calculation and hierarchical clustering and were finally grouped in 6 highly populated structural clusters (i.e. containing more than 10 molecules each) with distinct chemical scaffolds (Figure 3). Finally, out of these clusters, based on visual inspection of docking poses, logP and vendor availability, we shortlisted for experimental validation by NMR spectroscopy 21 compounds representative of the 6 clusters (Table 1).

**Figure 3.**
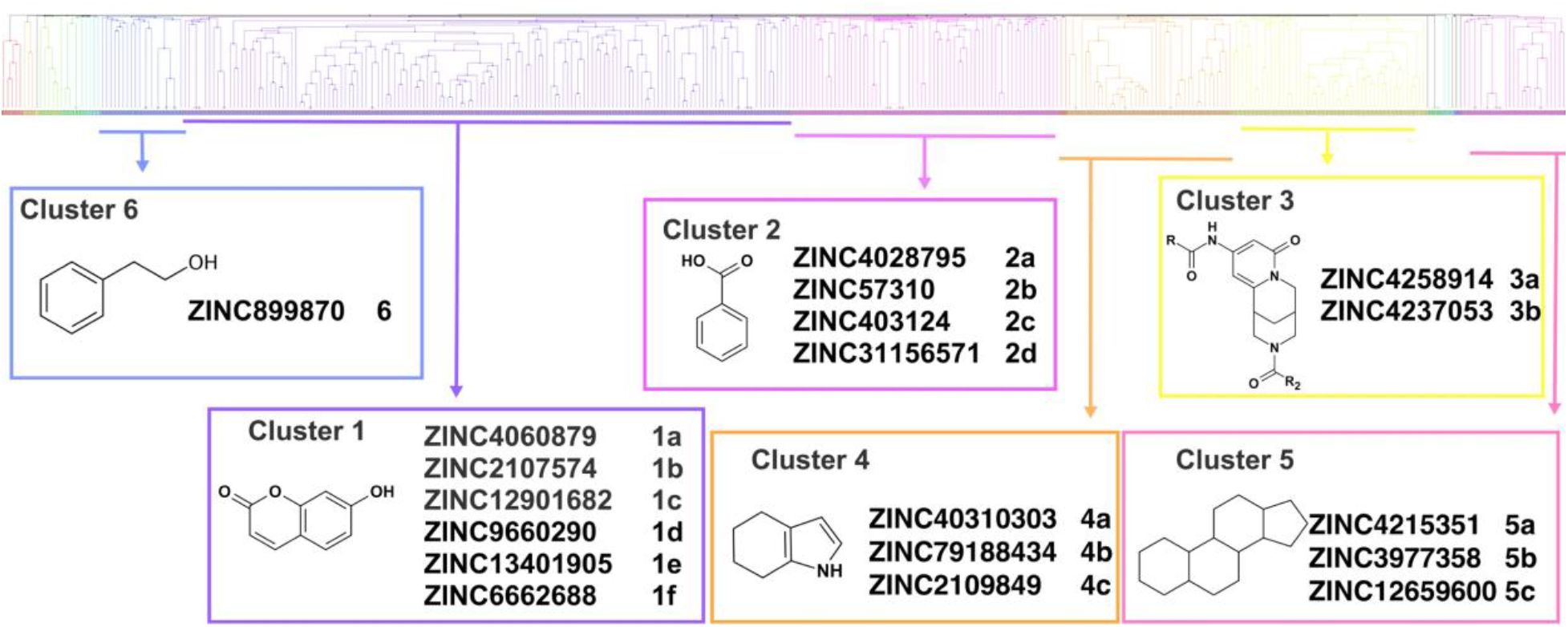
Cluster analysis. Dendrogram (obtained with Canvas 3.9) representing the hierarchical clustering based on chemical 2D fingerprint. 20 clusters were obtained, only those with a compound population with more than 10 molecules were considered, yielding 6 main clusters: cluster 1 (148 molecules, violet), cluster 2 (65 molecules magenta), cluster 3 (47 molecules, yellow), cluster 4 (40 molecules, orange), cluster 5 (25 molecules, pink), cluster 6 (20 molecules, light blue). For each cluster the chemical scaffold representative of the cluster, the ZINC code of the selected molecules for experimental validation and their corresponding IDs are reported.

**Table 1.**
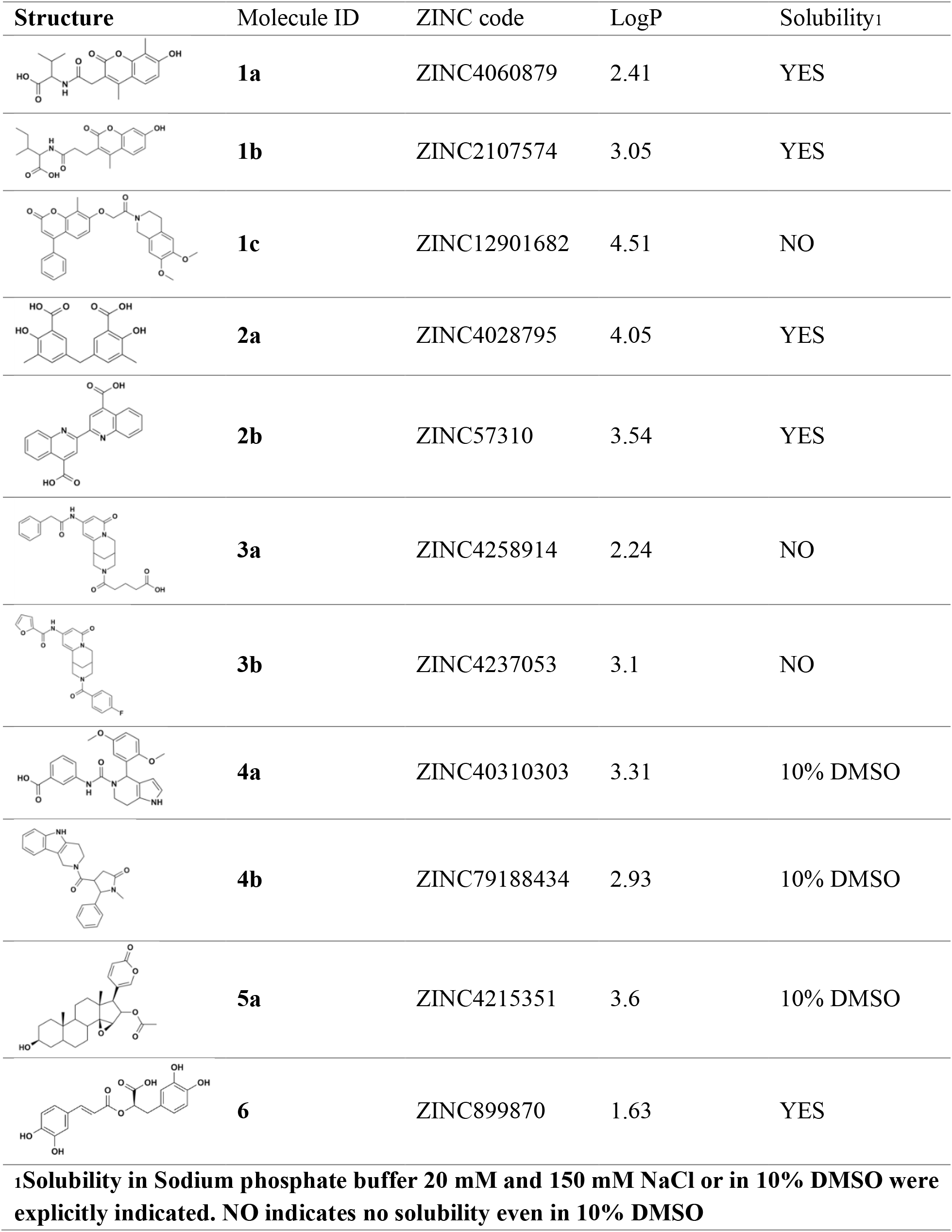
List of selected hits for experimental validation

| Structure | Molecule ID | ZINC code | LogP | Solubility<sub>1</sub> |
| --- | --- | --- | --- | --- |
|  | **1a** | ZINC4060879 | 2.41 | YES |
|  | **1b** | ZINC2107574 | 3.05 | YES |
|  | **1c** | ZINC12901682 | 4.51 | NO |
|  | **2a** | ZINC4028795 | 4.05 | YES |
|  | **2b** | ZINC57310 | 3.54 | YES |
|  | **3a** | ZINC4258914 | 2.24 | NO |
|  | **3b** | ZINC4237053 | 3.1 | NO |
|  | **4a** | ZINC40310303 | 3.31 | 10% DMSO |
|  | **4b** | ZINC79188434 | 2.93 | 10% DMSO |
|  | **5a** | ZINC4215351 | 3.6 | 10% DMSO |
|  | **6** | ZINC899870 | 1.63 | YES |
| **<sub>1</sub>Solubility in Sodium phosphate buffer 20 mM and 150 mM NaCl or in 10% DMSO were explicitly indicated. NO indicates no solubility even in 10% DMSO** |  |  |  |  |

### NMR validation of VS hits identifies 5,5’-methylenedi-2,3-cresotic acid (2a) as a ligand of HMGB1

Of the 21 compounds selected for experimental validation only 8 turned out to be soluble (> 0.1 mM) in water or in maximum 10% DMSO (Table 1). The binding to HMGB1 of the soluble molecules was next tested using ligand-based NMR spectroscopy methods. To this aim we prepared samples containing 50 μM HMGB1 and a 10-fold excess of ligand on which we performed saturation transfer difference (STD)^58^ and water-ligand observed via gradient spectroscopy (waterLOGSY) experiments^59^ (Figure 4 and Supplementary Figure S5). Of the compounds tested only **2a** (5,5’-methylenedi-2,3-cresotic acid) (Figure 4A) and **6** (rosmarinic acid) (Figure 4B) appeared to interact with HMGB1 as shown by STD effects and inversion of the sign in waterLOGSY experiments, whereby the strongest effects were observed for **2a** (Figure 4A).

**Figure 4.**
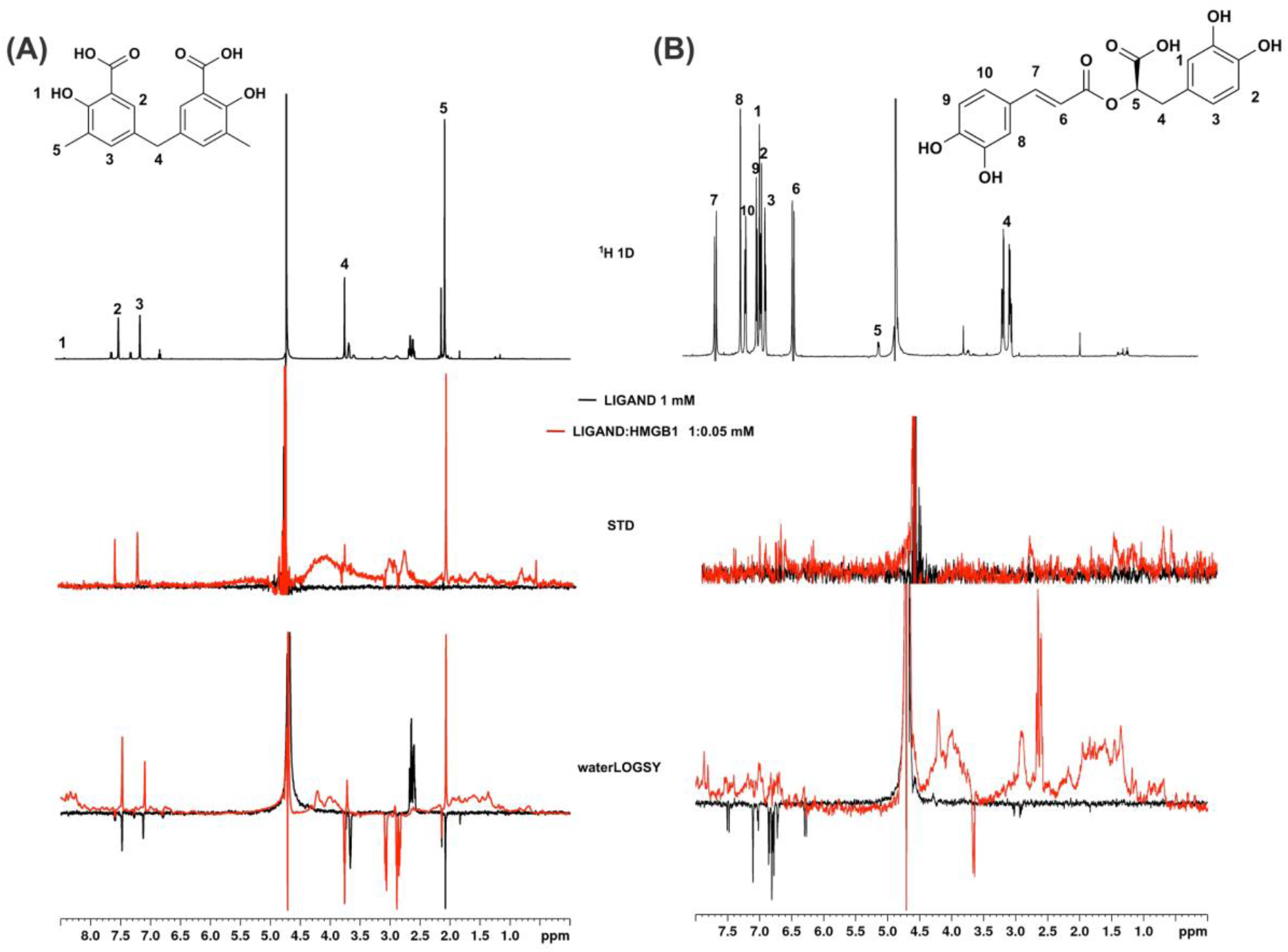
Ligand-based NMR binding assays performed on molecules 2a and 6 in the presence of HMGB1. For **2a** (A) and **6** (B) are represented: the chemical structure and the assigned ^1^H spectrum (top), the overlays of the STD spectra at 3s saturation (middle) and the waterLOGSY spectra (bottom). Experiments were performed on the ligand alone (1 mM) (black line) and in the presence of 0.05 mM HMGB1 (red line).

We further validated the interaction of these two ligands by protein-based NMR experiments^60^ using ^1^H-^15^N labelled HMGB1. Indeed, upon stepwise addition of **6** or **2a** we observed chemical shift perturbations (CSPs) in the HMGB1 Heteronuclear Single Quantum Coherence (HSQC) spectra (Figure 5A,B).

**Figure 5.**
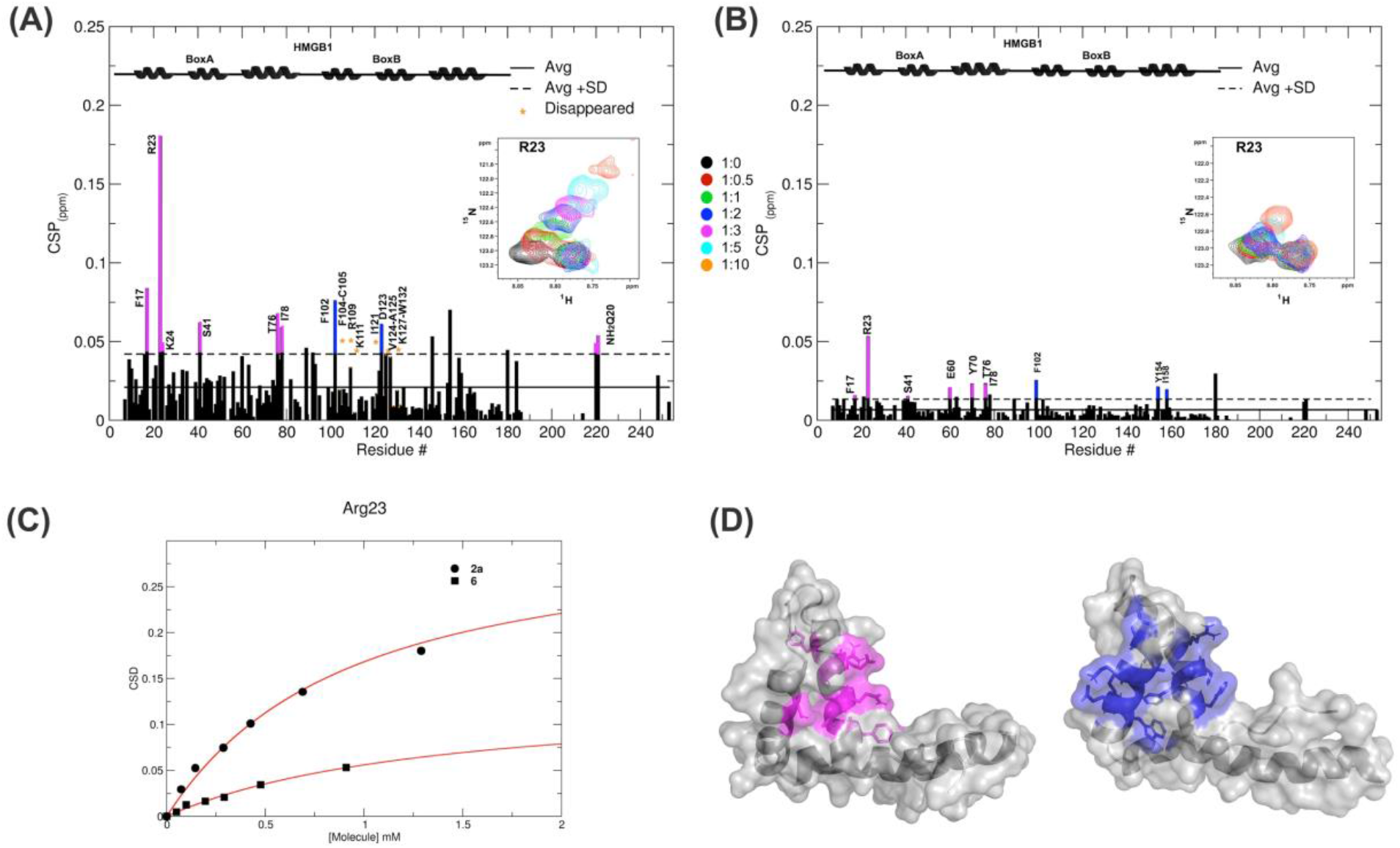
Interactions of 2a and 6 with HMGB1. Histogram showing the CSPs of ^15^N-labeled HMGB1 amides (~0.1 mM) upon addition of 10-fold excess of **2a** (A) and **6** (B). Missing residues are prolines or amides exchanging with the solvent, red stars indicate residues disappearing upon ligand binding. Elements of secondary structure are depicted on the top. BoxA and BoxB residues with CSP > Avg + SD are represented in magenta and blue, respectively. Inset: selected region of the superimposition of ^1^H-^15^N HSQC spectra of HMGB1 during the titration with the molecules (0.5, 1, 2, 3, 5 and 10 equivalents) showing the displacement of the peak associated to R23 during the titration. (C) Weighted average of R23 amide ^1^H and ^15^N chemical-shift changes as a function of increasing concentration of **2a** (circle) and **6** (square). Nonlinear curve fitting yields K_d_ = 0.9 ± 0.1 mM and >10 mM, respectively. (D) In grey surface and cartoon representations of BoxA and BoxB 3D structures, residues showing significant CSP (CSP > Avg + SD) upon **2a** binding are mapped in magenta and blue, respectively.

For molecule **2a** the interaction occurred on the fast-intermediate exchange regime on the NMR time scale with the disappearance of few peaks on BoxB (Figure 5A). On the other hand, the CSPs induced by **6** were in fast exchange regime and were by far smaller as compared to **2a**, and mainly involved BoxA (Figure 5B). Linear fitting of the chemical shifts as a function of added ligand indicated an apparent K_d_ of 0.9 mM and >10 mM for **2a** and **6**, respectively (Figure 5C). Interestingly, **6**, a polyphenolic component of the leaves of Perilla frutescens with general antinflammatory properties^61^, is known to attenuate inflammatory responses elicited by HMGB1/TLR4/NF-κB signaling^62,63^. This axis relies on the oxidized form of HMGB1, where Cys^22^ and Cys^44^ located on BoxA form a disulphide bond. On the other hand, the pattern of chemical shift perturbations (CSPs) (Figure 5A) clearly indicated that **2a** recognized both HMG boxes in full-length HMGB1. Similarly to both glycyrrhizin^26^ and diflunisal^24^, **2a** bound independently the two domains, and the pattern of CSPs observed for the isolated HMG boxes was comparable to the one in the full length protein (Supplementary Figure S6A-B). When mapped on the structures of the two HMG boxes, HMGB1 residues with significant CSP upon **2a** addition (F17, Q20, R23, E24, S41 on BoxA, F102, F104, C105, R109, K111, I121, D123, V124, A125, K127, L128, G129, E130, M131 and W132 on BoxB) defined a small surface between the first and the second helix (Figure 5D), in agreement with the computational analysis that identified this area as a putative hotspot (Figure 1). Indeed, this region on both boxes is characterized by a small solvent-exposed hydrophobic surface well suited for favorable van der Waals (vdW) interactions with the aromatic rings of **2a** (Figure 5D).

### 3D model of the interactions of 2a with HMGB1 boxes

CSPs data were next used to generate HADDOCK^49^ data-driven docking models of **2a** in complex with BoxA and BoxB. **2a** accommodates between the two short helices of the HMG boxes establishing favorable vdW interactions with the hydrophobic side chains of V19 and F37 in BoxA and of F102, V124 and A125 in BoxB (Figure 6A,B). For each box, the models suggest also the presence of stabilizing electrostatic interactions between one carboxylate of **2a** and the R23 and R109 sidechains of BoxA and BoxB, respectively (Figure 6A,B).

**Figure 6.**
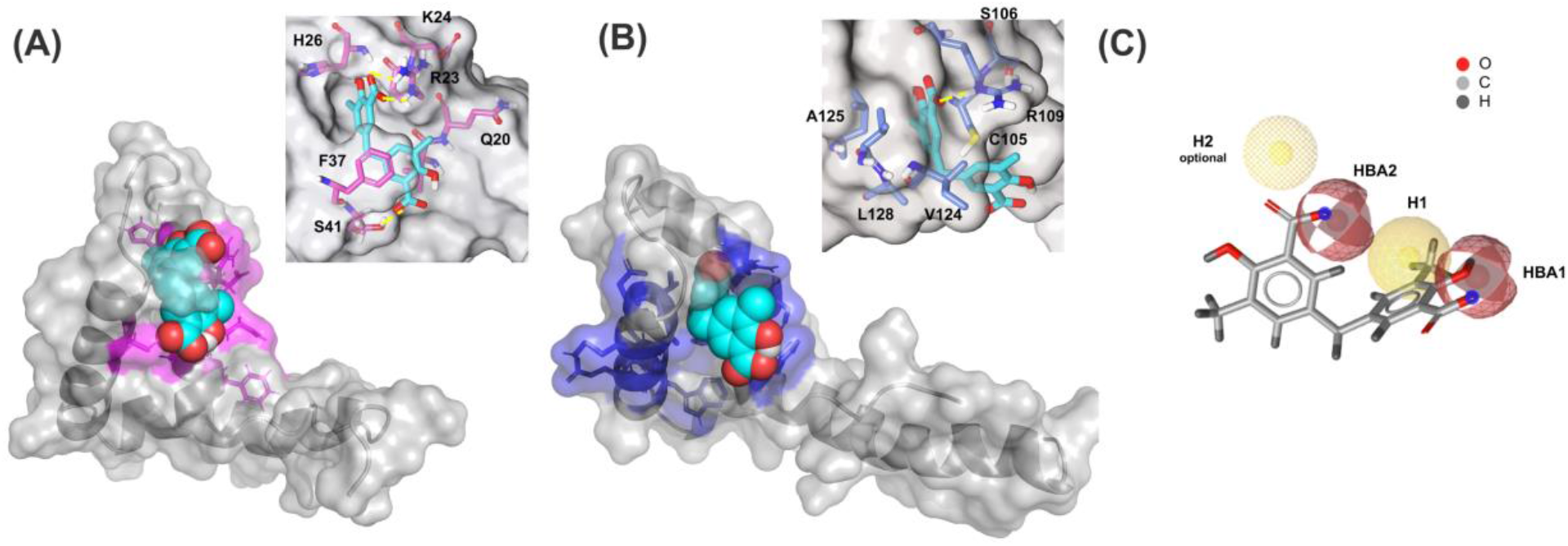
Three-dimensional models of 2a in complex with BoxA and BoxB. HADDOCK models of the interaction of **2a** (CPK representation) with BoxA (A) and Box B (B), residues with CSP > Avg + SD are colored in magenta and blue, respectively. In the insets HMGB1 residues (sticks) involved in hydrophobic and electrostatic interactions with **2a** are explicitly labelled. Fitting of **2a** structure with PhMOD. **2a** fulfils 3 out 4 pharmacophoric features, consisting in two hydrophobic (H1 and H2, the latter being optional) and two H-Bonding acceptor features (HBA1 and HBA2), shown in yellow and red, respectively.

Indeed, in NMR titrations performed on a ^15^N-HMGB1 mutant, where both R23 and R109 were substituted by alanines (R23A/R109A), we observed reduced chemical shift displacements, indicative of reduced interaction, thus supporting the involvement of these arginines in binding to **2a** (Supplementary Figure S7). The second aromatic ring and the associated carboxylic group of **2a** establish π-π and polar interactions with F37 and S41, respectively (Figure 6A). Conversely, in BoxB the second salicylate moiety does not seem to be involved in stabilizing interactions (Figure 6B). Collectively, these 3D models indicate that the steric and electronic features of **2a**, which fulfill the predicted pharmacophoric model (Figure 6C), are indeed appropriate to interact with both HMG boxes. In particular, the two major interactions between the ligand and the target consist in *i*. a salt-bridge between the carboxylate of **2a** and the guanidinium groups of the conserved R23 and R109, and *ii*. hydrophobic interactions between the phenyl groups of **2a** and the hydrophobic patch at the interface of the two helices forming the short arm of the L shaped HMG boxes.

### 2a breaks the HMGB1•CXCL12 heterocomplex

NMR titrations and HADDOCK calculations indicate that **2a** targets in part the region that has been shown to be involved in the interaction with CXCL12^24,25,64^. We therefore asked whether **2a** was able to interfere with the HMGB1•CXCL12 heterocomplex (Figure 7A). To this end we performed MST experiments, whereby increasing concentrations of **2a** were added to a preformed HMGB1•CXCL12 complex. During the titration we observed a sharp transition at 0.31 ± 0.04 mM, consistent with the detachment of fluorescently labelled HMGB1 from the heterocomplex (Figure 7A). These results were further confirmed by NMR-based Antagonist Induced Dissociation Assay (AIDA)^45^. In this experiment a ^15^N HSQC spectrum was first acquired on free ^15^N HMGB1 (Figure 7B), then on a preformed complex composed by ^15^N labelled HMGB1 (0.1 mM) and unlabeled CXCL12 (0.2 mM). The spectrum appearance was characterized by broadening of HMGB1 resonances (Figure 7C), ascribable to the intermediate regime on the NMR time scale and to the increased relaxation time upon complex formation^24^. Addition of **2a** (0.2 mM) to the preformed ^15^N-HMGB1•CXCL12 heterocomplex resulted in a drastic line broadening of the ^1^H-^15^N HSQC spectrum that caused the disappearance of the majority of HMGB1 peaks (Figure 7D). We interpreted this strong line broadening as an effect associated to multiple equilibria involving free HMGB1, HMGB1 bound to **2a**, HMGB1 bound to CXCL12. Upon addition of ten-fold excess of **2a** we observed the narrowing of ^1^H-^15^N signals providing direct evidence of complex dissociation (Figure 7E). Taken together both MST and AIDA experiments indicate that **2a** impairs the HMGB1•CXCL12 heterocomplex.

**Figure 7.**
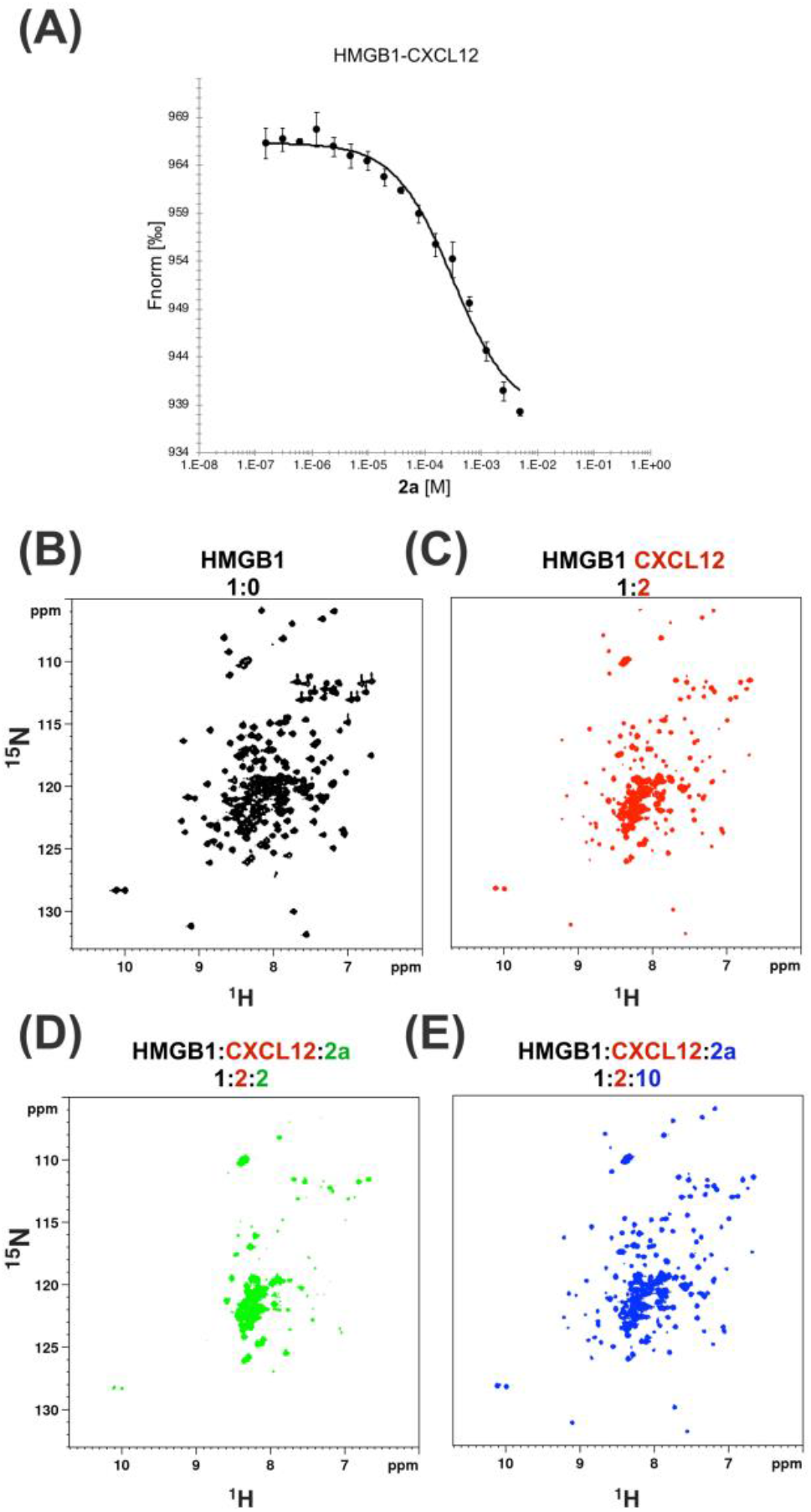
2a breaks HMGB1•CXCL12 heterocomplex. MST curve of HMGB1•CXCL12 heterocomplex (preformed using 50 nM HMGB1 and 10 □ M CXCL12) in the presence of increasing concentrations of **2a**. Decreasing MST signal (from ~965 to ~935 a.u.) upon **2a** addition indicates that CXCL12 has been displaced from HMGB1. n = 3; data represent Avg ± SD. ^1^H-^15^N HSQC HMGB1 (0.1 mM) spectrum (B) without (black), (C) with 0.2 mM CXCL12 and upon addition of (D) 0.2 mM and (E) 1 mM of **2a**.

### 2a binds to CXCL12

Recent work on the inhibition of HMGB1 chemotactic activity by diflunisal has revealed a peculiar mechanism of action in which the breakage of HMGB1•CXCL12 interaction occurs through the dual binding of the ligand to both proteins^24^. In the same study we also demonstrated that glycyrrhizin, the first chemical probe identified to bind and inhibit HMGB1 chemotactic activity^26^, was a binder of CXCL12 and an inhibitor of the HMGB1•CXCL12 heterocomplex. We thus asked whether **2a** was also able to bind CXCL12. Indeed, NMR titrations of **2a** on ^15^N labelled CXCL12 confirmed a direct interaction (Figure 8A). Binding occurred in the fast exchange regime, in line with the 200 μM affinity, as measured by MST (Supplementary Figure S8A) and by NMR (Supplementary Figure S8B). The highest CSPs upon ligand binding involved residues located on the β1 strand (V23, H25, K27) and on the so called CXCR4 sulfotyrosine (sulfoY21) binding site^65^ (N45, Q48, V49) (Figure 8A). Indeed STD and waterLOGSY competition experiments performed adding 1 mM **2a** to a preformed CXCL12:diflunisal complex (1:10) indicated that **2a** competes with diflunisal for the same binding site, as assessed by the reduction of both STD and waterLOGSY signals of diflunisal upon **2a** addition (Figure 8B).

**Figure 8.**
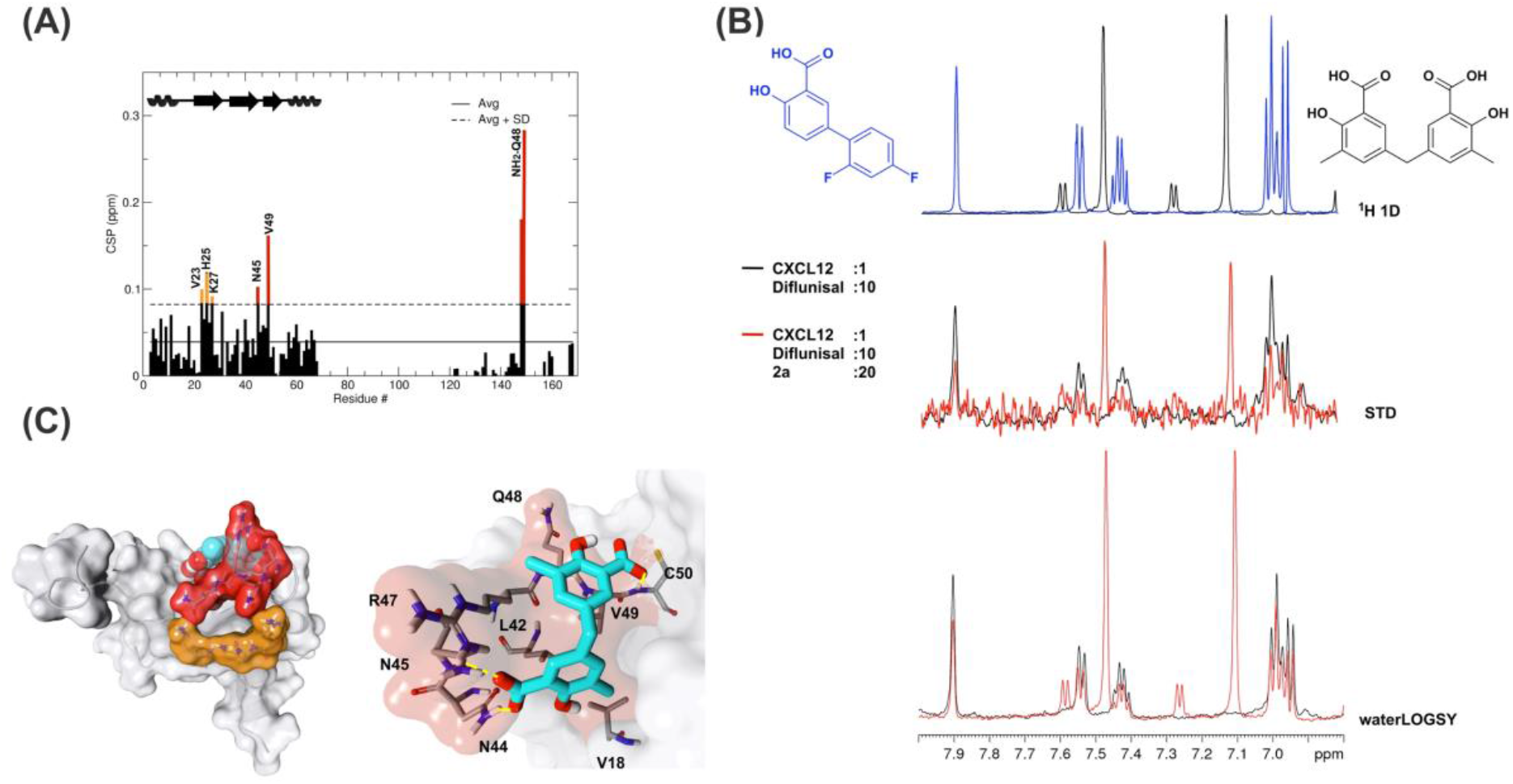
2a binds to CXCL12. **(A)** Histogram showing the CSPs of ^15^N-labeled CXCL12 amides (~0.1 mM) upon addition of 10-fold excess of **2a**. Missing residues are prolines. Elements of secondary structure are depicted on the top. (B) Top: ^1^H spectra of **2a** (black) and of diflunisal (blue). Middle: superimposition of STD spectra (3s saturation) obtained on 0.5 mM Diflunisal with 0.05 mM CXCL12 (black line) and upon addition of 1 mM of **2a** (red line). Bottom: waterLOGSY spectra obtained for 0.5 mM diflunisal with 0.05 mM CXCL12 (black line) and upon addition of 1 mM of **2a** (red line). The decrease of signal intensities of diflunisal upon **2a** addition indicates that the two ligands compete for the same binding site (C) Left: HADDOCK model of the interaction of **2a** (CPK representation) with CXCL12 (grey surface and cartoon). CXCL12 residues with CSP > Avg + SD located around the sY21 binding site on the β1 strand are represented in red and orange, respectively. Right: Zoom in of the binding site, CXCL12 residues (sticks) involved in hydrophobic and polar interactions with **2a** (sticks) are explicitly labelled.

As for diflunisal, we hypothesized that CSPs affecting residues of the β1 strand were due to allosteric effects induced by ligand binding. A HADDOCK model of the **2a**-CXCL12 complex indicated that **2a**, in analogy to diflunisal^24^, accommodates in the CXCR4 sulfoY21 binding site, whereby the two salicylate moieties of **2a** establish polar interactions with R47, N44 and N45 sidechains and with the backbone carbonyl of Q48 and the amide of C50, respectively. Both aromatic rings are involved in hydrophobic interactions with V18, V49, L42 (Figure 8C). Taken together these data show that **2a** is also a ligand of CXCL12 and targets the CXCR4 sulfoY21 binding site.

### 2a inhibits the chemotactic activity elicited by HMGB1•CXCL12 heterocomplex

Finally, having assessed the ability of **2a** to simultaneously bind HMGB1 and CXCL12 and to break the HMGB1•CXCL12 heterocomplex, we asked whether **2a** affected the chemotactic activity of the HMGB1•CXCL12 heterocomplex. Indeed, **2a** inhibited the chemotaxis of mouse 3T3 fibroblasts in a dose-dependent manner, with an IC_50_ close to 10 pM, the lowest documented so far for inhibitors of HMGB1–induced cell migration. Importantly, **2a** did not influence the general motility of fibroblasts, as it did not affect chemotaxis toward fMLP (Figure 9), indicating that **2a** targets selectively the chemotactic activity of the HMGB1•CXCL12 heterocomplex.

**Figure 9.**
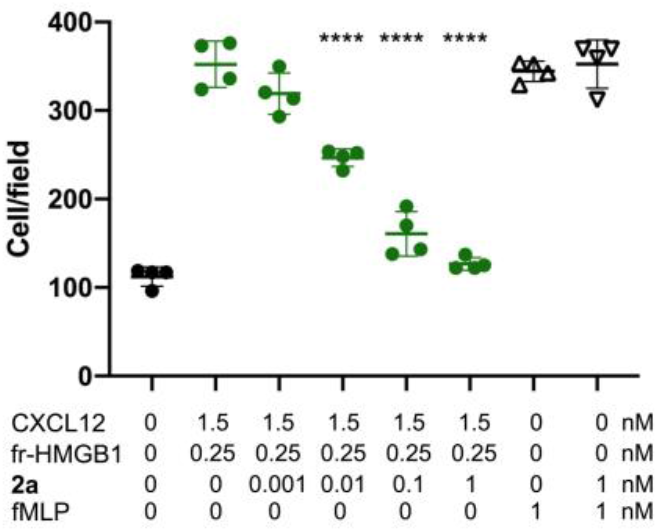
2a inhibits HMGB1•CXCL12 heterocomplex-induced chemotaxis, Mouse 3T3 fibroblasts were subjected to chemotaxis assays in Boyden chambers, 0.25 nM HMGB1, 1.5 nM of CXCL12 or no chemoattractant was added in the lower chamber, together with the indicated concentrations of **2a**. **2a** does not inhibit fMLP-induced chemotaxis at the highest concentrations tested for HMGB1•CXCL12 induced chemotaxis. Data points with average ± standard deviation (Avg ± SD; n = 3, each point represents a biological replicate) in a representative experiment. Statistics: one-way ANOVA (P < 0.0001), followed by Dunnett’s post-tests. ****P < 0.0001 relative to no **2a** addition.

## Discussion

Recent studies investigating the functional synergism of chemokine-based heterodimers have shown that hampering heterophilic interactions interrupts inflammation^8^. Thus, the targeting of these PPIs is emerging as a valuable strategy for the development of new selective antagonists suitable for the tailored modulation of specific inflammatory responses. In this sense a very promising example is represented by CCL5-derived peptides that inhibited the atherogenic CCL5•CCL17 interaction and hampered CXCL12-driven platelet activity^8^. However, the development of PPI antagonists, either peptidomimetics or small organic molecules, poses incredible challenges when the three-dimensional structure of the complex is unknown. This is the case for HMGB1•CXCL12 heterocomplex, a master-regulator of the recruitment of inflammatory cells via the CXCR4 axis, and therefore a valuable target for the development of selective anti-inflammatory compounds. However, high-resolution atomistic descriptions of this interaction are still elusive, thus making structure-based drug discovery studies extremely difficult. One reductionist strategy that can be adopted to cope with this problem is to target the isolated components of the heterocomplex. While CXCL12 as direct interactor of CXCR4 is a popular target for drug development, as shown by several structure-based drug designs studies^29,66^, no systematic structure-based small-molecule screening has been reported until now for HMGB1. This is probably due to the structural features of both the HMG boxes, that do not present clearly targetable pockets. Still, computational analysis of their structures through the mapping servers FTMap^32^ and Dogsite^34^ performed in this work highlighted a binding hot spot at the interface of helix I and helix II of both HMG boxes. Interestingly, the same region is recognized by several known HMGB1 binders, including DNA^67^, small ligands glycyrrhizin^26,38^ and diflunisal^24^), peptides^68^ or proteins such as p53^69^, speaking in favor of virtual screening approaches targeting this area. Prompted by these results we screened the Zbc library with three different ligand docking programs that rely on different search algorithms and scoring functions, with the aim to compensate for their individual weaknesses^55,56^. The ligands of the best docking poses emerging from these three docking programs were *i*. scrutinized and filtered according to a pharmacophoric model based on known HMGB1 binders, *ii*. clustered according to structural similarity and *iii*. subsequently validated through ligand-based and protein-based NMR methods. Interestingly, at the end of this pipeline rosmarinic acid (**6**) emerged as a weak ligand of HMGB1. Indeed, this molecule has been already reported to be an attenuator of the HMGB1/TLR4/NF-κB-dependent inflammatory response ^62,63^, supporting the validity of our virtual screening approach. We then focused on the novel hit stemming from our pipeline, 5,5’-methylenedi-2,3-cresotic acid (**2a**). **2a** was previously reported to have modest DNA demethylase inhibition activity^70^. We demonstrated that **2a** interacts with high micromolar affinity with the predicted hotspot of the single boxes in a way similar to what observed with diflunisal, targeting the surface directly involved in the formation of the heterocomplex with CXCL12^24^. Accordingly, NMR and MST competition experiments demonstrated that **2a** is able to disrupt the interaction of HMGB1 with CXCL12, with an EC_50_ of 0.3 mM. Moreover, cell migration experiments showed that **2a** is able to inhibit the chemotactic activity of HMGB1•CXCL12 heterocomplex, with an IC_50_ of 10 pM, the most potent inhibition activity documented until now, and 2 orders of magnitude larger than that of diflunisal. Intriguingly, NMR CSP and MST data indicate that **2a** binds also to CXCL12, as does diflunisal. It targets with high micromolar affinity the CXCR4 sulfotyrosine binding pocket and induces CSPs on CXCL12 β1 strand, likely ascribable to allosteric effects^71^ contributing to the destabilization of the interaction with HMGB1. The dual binding ability to HMGB1 and CXCL12 seems to be a common theme in inhibitors of HMGB1•CXCL12 chemotactic activity, as observed for glycyrrhizin and diflunisal^24^. Results emerging from this and previous studies strongly suggest that salicylate derivatives are well suited for the interaction with HMGB1, with the carboxylic and hydroxylic group of the aromatic rings fulfilling the predicted pharmacophoric characteristics required for HMGB1 ligands. Intriguingly, these pharmacophoric features seem in part to satisfy also the chemical requirements for CXCL12 ligands. In fact, previous structure-based design studies aiming at identifying CXCL12 inhibitors have suggested that a carboxylic group, preferentially combined to an aromatic group, is a good mimic of the negatively charged CXCR4 sulfotyrosine^29,66^. Thus, we speculate that some known CXCL12 inhibitors might be also ligands of HMGB1, conceivably exerting a synergic inhibition effect on the disruption of the HMGB1•CXCL12 heterocomplex and as direct inhibitors the interaction of CXCL12 with CXCR4.

The double targeting action of **2a**, diflunisal and glycyrrhizin only in part reconciles the discrepancy existing between their high micromolar ability to disrupt the HMGB1•CXCL12 complex and their nanomolar or picomolar chemotactic inhibition efficacy. Chemotaxis relies on many different signaling cascades at the cell membrane, where the local concentration of both HMGB1 and CXCL12 might increase through their direct interaction with cell surface glycosaminoglycans (GAGs)^72,73^. Additional mechanisms and cooperative binding phenomena among multiple actors at the cell surface might occur^74^ herewith synergistically and/or allosterically contributing to complex destabilization. We posit that the inhibition activity of these ligands might rely on multistep dynamic processes that, besides the interaction with both CXCL12 and HMGB1, also include the direct binding to a high affinity binding pocket at the heterocomplex interface. Once targeted, this site might cooperate to heterocomplex dissociations and/or induce allosteric changes that affect heterocomplex binding to CXCR4 and the related signaling. As long as the three-dimensional structure of the heterocomplex is unknown and the molecular and functional details of its interaction with CXCR4 remain underexplored, the details of the mechanisms dictating the inhibitory potency of **2a** and of other ligands of the HMGB1/CXCL12/CXCR4 axis will be unresolved. Nevertheless, our reductionist approach showed that small solvent-exposed clefts of HMGB1 boxes, traditionally viewed as a barrier to the success of *in silico* ligand screening, can actually serve as legitimate sites for drug discovery. Even more interestingly, cytokine heterocomplexes that appeared undruggable are actually promising drug targets.

## Conclusions

In summary, the combination of VS and experimental validation proved successful in identifying **2a** as the best molecule documented until now in terms of inhibition of *in vitro* chemotaxis activity of the HMGB1•CXCL12 heterocomplex. These results pave the way for future structure activity relationship studies for the optimization of the pharmacological targeting of this heterophilic interaction for anti-inflammatory purposes.

## Author Contributions

F.D.L. performed the NMR and MST experiments, computational studies, analyzed the data and prepared the manuscript. G.Q. performed the NMR experiments, M.M. expressed and purified recombinant proteins. F.D.M. performed the cell migration experiments. M.E.B. and G.M directed the study and were involved in all aspects of the experimental design, data analysis and manuscript preparation. All authors critically reviewed the text and figures.

## Supplementary Material

Training set (TS), migration of decoys, building and validation of the Pharmacophore model (PhMOD), HADDOCK restraints, HADDOCK clusters, hits binding to HMGB1 by ligand-based NMR methods, Binding of 2a to BoxA and BoxB, comparison 2a binding to HMGB1 wt and R23A/R109A, MST and NMR binding curve of 2a to HMGB1.

