## Supplementary material for "Discovery of 5,5’-methylenedi-2,3-cresotic acid as a potent inhibitor of chemotactic activity of the HMGB1•CXCL12 heterocomplex using Virtual Screening and NMR validation": Supplementary Material.pdf

**Figure S1** *List of molecules used* as *Training set (TS) to generate the PhMOD*
**Figure S2** *Effect of decoy molecules on HMGB1 induced chemotaxis*
**Figure S3** *Definition of the Pharmacophore model (PhMOD).*
**Figure S4** HADDOCK clusters
**Figure S5** *Validation of hits by ligand-based NMR methods*.
**Figure S6** *Histograms of CSPs induced by* ***2a*** *on isolated BoxA and BoxB*
**Figure S7** *Comparison of the CSPs induced by* ***2a*** *binding to HMGB1 and to R23A/R109A* double *mutant*
**Figure S8** *MST and NMR isothermal binding curves of* ***2a*** *to HMGB1*
**Supplementary Table S1** *List of Ambiguous Interaction Restraints (AIRs) and of Unambiguous restraints (nOe restrains) used in HADDOCK calculations*.

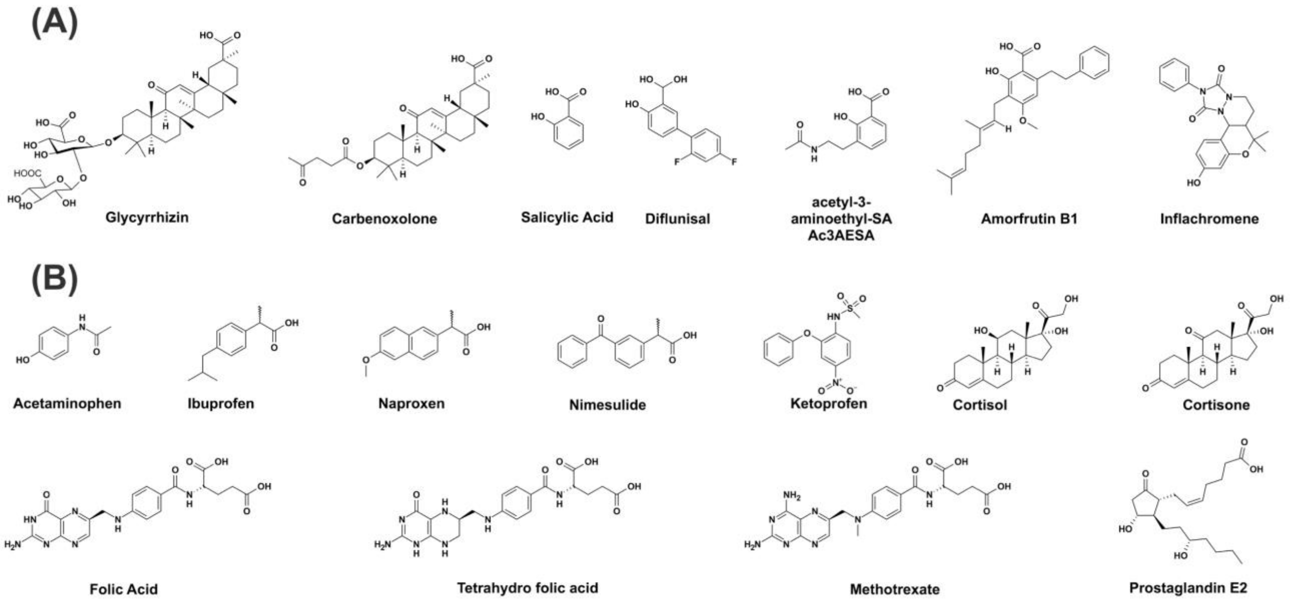


**Supplementary Figure S1**. *List of Molecules used as Training set (TS) to generate the PhMOD* Chemical structures of the 7 known active HMGB1 molecules (A) and of the 11 decoys (B) composing the TS.

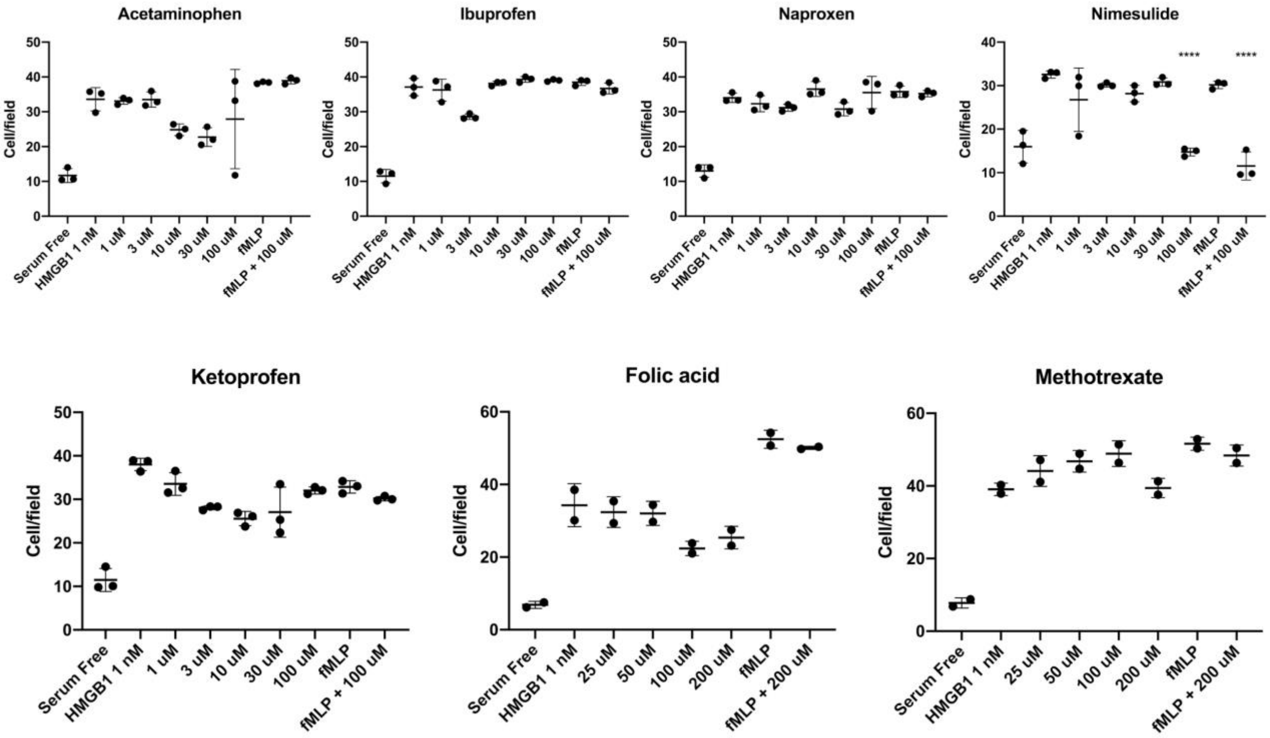


**Supplementary Figure S2**. *Effect of decoy molecules on HMGB1 induced chemotaxis.* Mouse 3T3 fibroblasts were subjected to chemotaxis assays in Boyden chambers, 1 nM HMGB1, or no chemoattractant was added in the lower chamber, together with the indicated concentrations of the molecules. Data points with average ± standard deviation (avg ± SD; n = 3 for acetaminophen, ibuprofen, naproxen, nimesulide and ketoprofen, n = 2 for Folic Acid and methotrexate each point represents a biological replicate) in a representative experiment. Statistics: one-way ANOVA ($P < 0.0001$), followed by Dunnett's post-tests. ****$P < 0.0001$ relative to no molecule addition. Decoys do not inhibit HMGB1-induced cell migration, or when they do so (100 µM nimesulide), they also inhibit fMLP-induced cell migration.

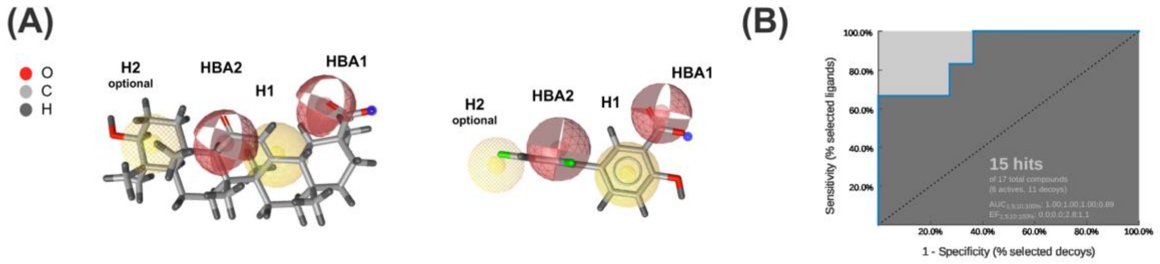


**Supplementary Figure S3**. *Definition of the Pharmacophore model (PhMOD).* (A) Fitting of Glycyrrhizin and diflunisal structures (represented in sticks) into the PhMOD. The PhMOD consists of two hydrophobic features (H1 and H2, the latter being optional) shown in yellow, and of two H-Bonding acceptor features (HBA1 and HBA2) shown in red. (B) ROC curve and AUC resulting from the evaluation of the training set: 15 hits of 18 queries (7 actives, 11 decoys) AUC=0.89. The ROC plot was calculated and visualized using LigandScout3.02.

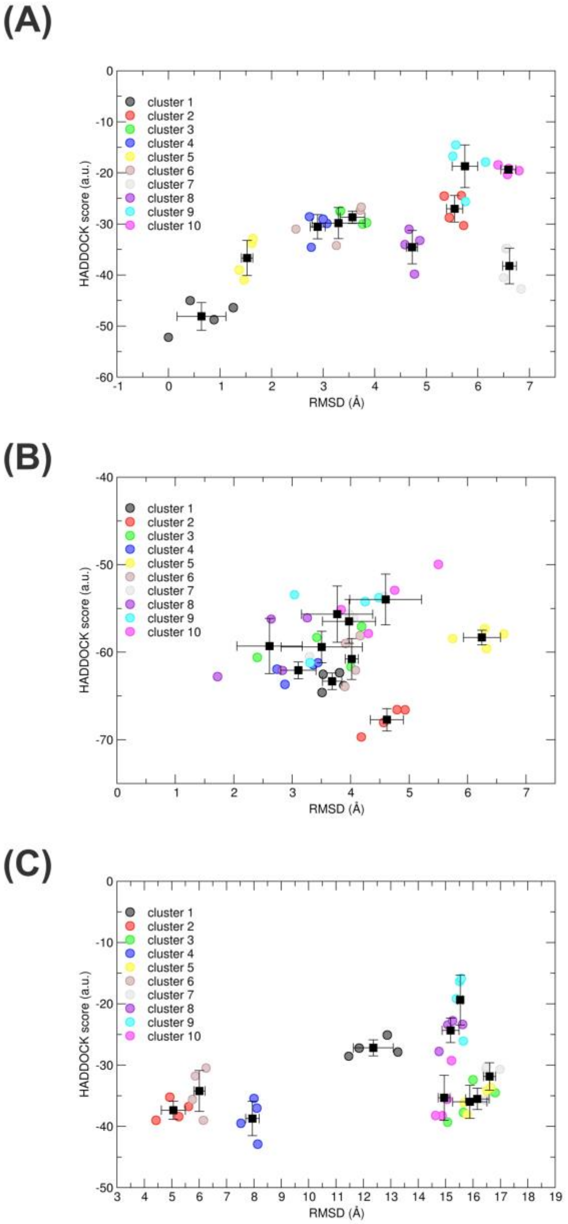


**Supplementary Figure S4**. *Data-driven molecular docking models*. HADDOCK score *versus* rmsd from the lowest Haddock energy complex structure between **2a** and BoxA (A), and BoxB (B) and CXCL12 (C). Circles correspond to the four best structures in each cluster, the cluster averages with the standard deviation are indicated with the black squares and bars.

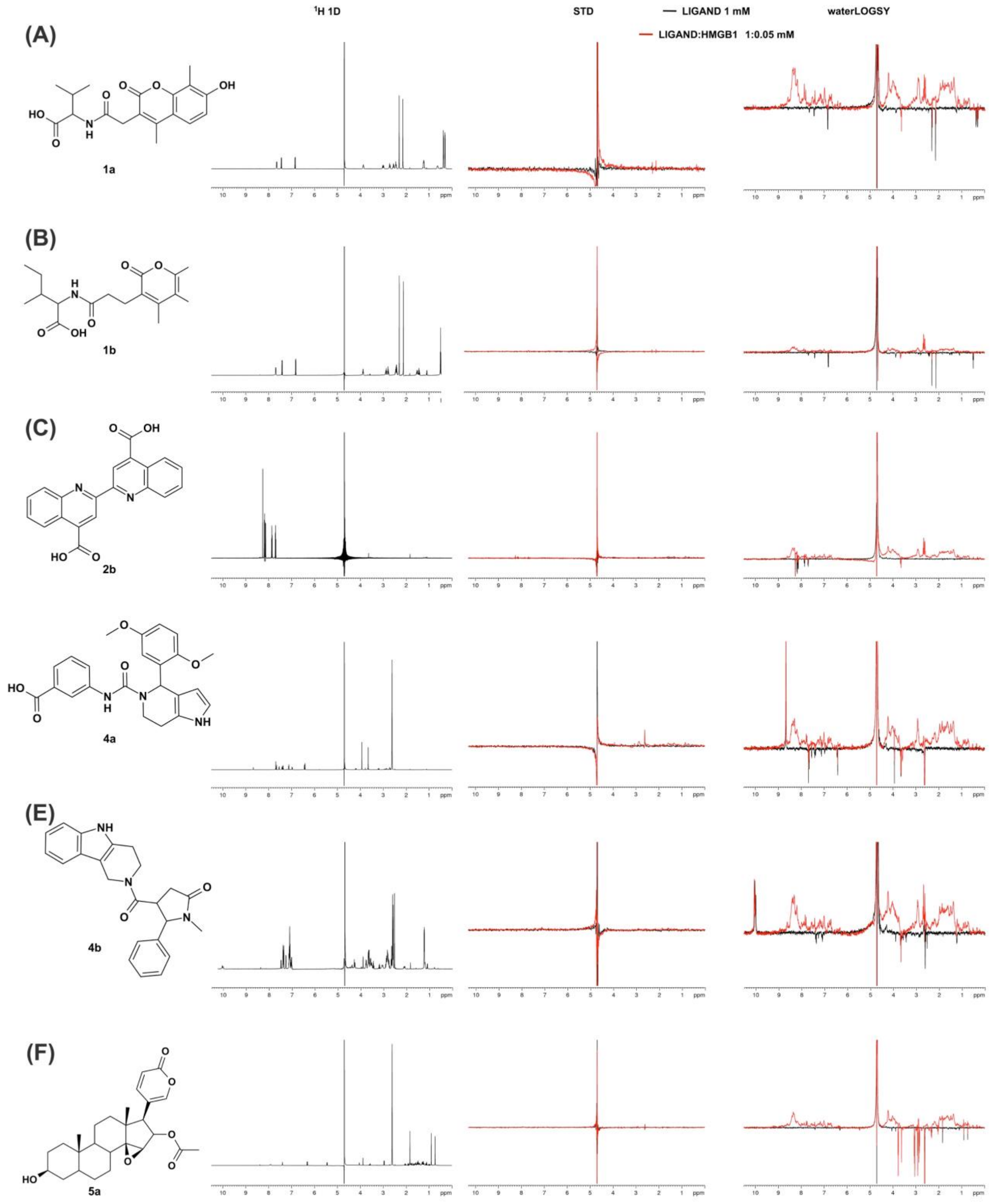


**Supplementary Figure S5**. *Validation of hits by ligand-based NMR methods*. Chemical structures, 1H spectra, overlay of STD spectra obtained for 1 mM of ligand alone (black line) and with 0.05 mM HMGB1 (red line), waterLOGSY spectra obtained for 1 mM of ligand alone (black line) and with 0.05 mM HMGB1 (red line) are shown for molecule **1a** (A), **1b** (B), **2b** (C), **4a** (D), **4b** (E) and **5a** (F). The five molecules do not interact with HMGB1. Molecules **1a**, **1b**, **4b** were also tested in protein-based NMR experiments with 15N-labelled HMGB1 and did not show any binding (data not shown).

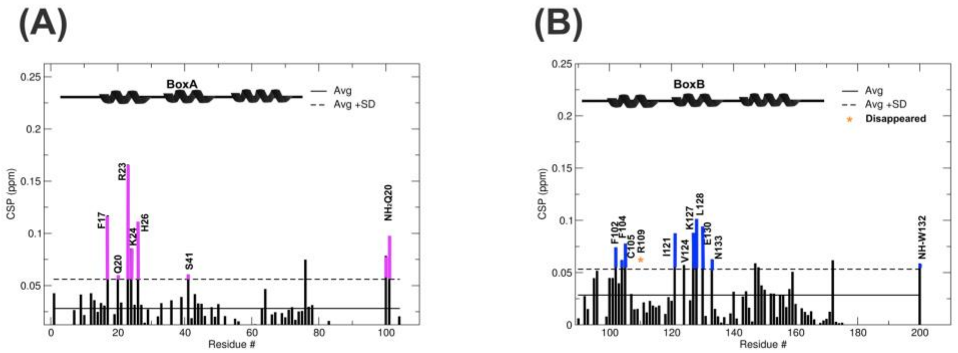


**Supplementary Figure S6.** *Histograms of CSPs induced by **2a** on BoxA and BoxB*. Histograms of CSPs (ppm) of (A) BoxA and (B) BoxB amide resonances upon addition of ten-fold excess of **2a**. CSPs > Avg + SD are highlighted in magenta (BoxA) and in blue (BoxB). α-helices are schematically represented on the top.

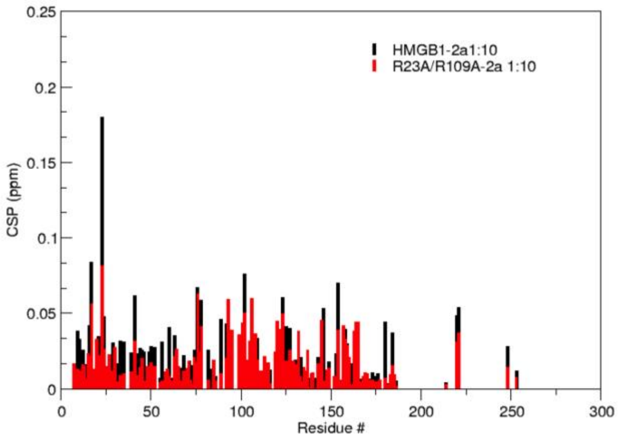


**Supplementary Figure S7.** *Comparison of the CSPs induced by **2a** binding to HMGB1 and to R23A/R109A double mutant*. Histograms of CSPs (ppm) of HMGB1 (black) and R23A/R109A double mutant (red) amide resonances upon addition of ten-fold excess of **2a**. The lower CSPs values induced by **2a** on R23A/R109A double mutant indicate a reduced interaction.

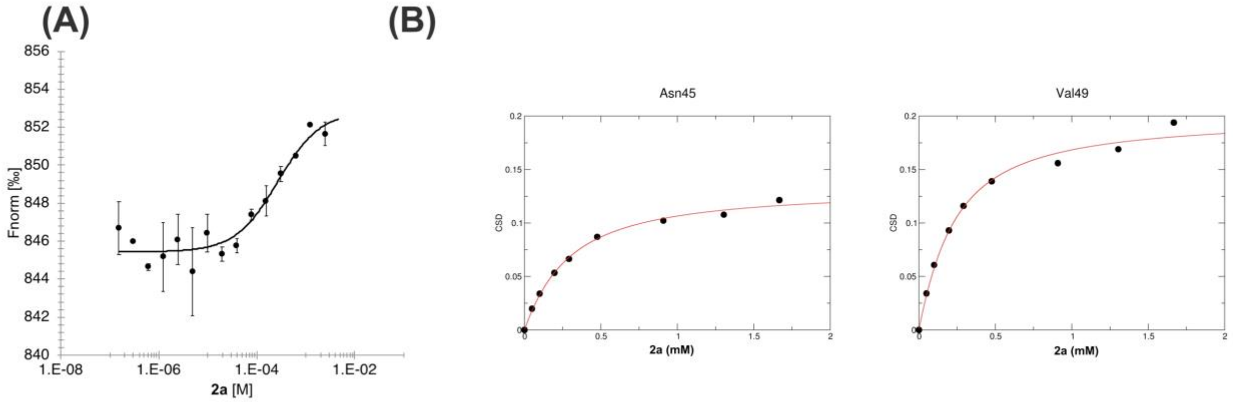


**Supplementary Figure S8.** *MST and NMR isothermal binding curves of **2a** to HMGB1*. (A) Dose-response curve of CXCL12-**2a** interaction followed by MST. **2a** concentrations ranged from 5 mM to 153 nM on 50 nM labelled 6His-CXCL12. The $K_d$ is 0.27 ± 0.1 mM (n=3, error bars correspond to sd) (Fnorm=normalized fluorescence). (B) Weighted average of N45 and V49 amide $^{1}H$ and $^{15}N$ chemical-shift changes in the presence of increasing concentration of **2a**. Nonlinear curve fitting yields $K_d$ = 200 ± 25 μM.

**Supplementary Table S1.** List of Ambiguous Interaction Restraints (AIRs) and of Unambiguous (nOe) restraints used in HADDOCK calculations.

| | Ambiguous | | Unambiguous[a,b] |
|---|---|---|---|
| Domain | Active | Passive | |
| Box A | F17, V19[nOe], Q20, R23, E24, S41 | Y15, F17, F18, V19, Q20, R23, E24, H26, K27, V35, F37, E40, S41, C44, W48 | HG*$_{V19}$-H3 $_{\mathbf{2a}}$; or HG*$_{V19}$-H5 $_{\mathbf{2a}}$ |
| Box B | P102, P104, C105, T108, R109, I121, V124, A125, L128 | L103, S106, E107, P110, K111, K113, G122, D123, K127, G129, E130, M131, W132 | *n.d.* |
| CXCL12 | E15, V18, N45, R47 Q48, V49 | C11, V23, K24, H25, L26, K27, Q48 | *n.d.* |

*n.d.* Not determined
[a] OPLS *force field* nomenclature of protein hydrogens
[b] **2a** hydrogens (Figure 4A)
